# Neural Dynamics of Object Manifold Alignment in the Ventral Stream

**DOI:** 10.1101/2024.06.20.596072

**Authors:** Binxu Wang, Carlos R. Ponce

**Affiliations:** Department of Neuroscience, Washington University in St Louis 660 S. Euclid Ave., St Louis, 63110, MO, USA; Department of Neurobiology, Harvard Medical School 220 Longwood Avenue, Boston, 02115, MA, USA

**Keywords:** GAN, deep network, Ventral Stream, neural code

## Abstract

Visual neurons respond across a vast landscape of images, comprising objects, textures, and places. Natural images can be parameterized using deep generative networks, raising the question of whether latent factors learned by some networks control images in ways that better align with visual neurons. We studied neurons in areas V1, V4 and posterior IT, optimizing images using a closed-loop evolutionary algorithm. We used two generative image models: (1) DeePSim, which parameterizes local image patterns, and (2) BigGAN which parameterizes object identity and nuisance variables. We found that neurons could guide image optimization on both pattern- and object-based image manifolds across areas; V1 aligned best with the DeePSim image space, whereas PIT aligned well with both DeePSim and BigGAN spaces. While initially PIT neurons responded well to the textural manifold, their responses to objects also emerged over time, suggesting that object-like responses required further processing. We identified similar local features common to both textural and object images, but not optimal global configuration. We conclude that visual cortex neurons are aligned to a representational space not yet captured by current artificial model of the visual system.

---

To understand the visual brain, we must examine what activates its constituent neurons. Highly activating images provide information about the latent factors behind the tuning of these neurons. With the advance of generative image models, deep neural networks can be trained to generate natural images from sampled latent code, effectively parameterizing the space of natural images [1, 2]. As such, they allow for the systematic study of neuronal selectivity. Using generative models in combination with evolutionary algorithms, we have previously synthesized highly activating images for neurons across the ventral stream in the primate[3], finding intriguing trends across the visual hierarchy [4, 5, 6]. These studies relied on one specific image model, a generative network (GAN) called DeePSim [7]. The rapid progress of generative models provides an opportunity to test the generality of both our method and results. We leveraged our understanding of the geometry of a generative network’s latent space[8] and their most suitable evolutionary algorithms[9], to apply the neural-guided image synthesis paradigm to more recent image generators. We compared the influence of different generative manifolds on the visual feature preferences of individual neurons in V1, V4, and posterior inferotemporal cortex (PIT). Specifically, we compared neuron-guided image synthesis in two generative adversarial networks (GANs): the older but powerfully expressive DeePSim GAN [7] and the more modern and photo-realistic BigGAN [10]. DeePSim has served as an effective natural image prior for previous vision research [3, 4, 5], but it does not generate whole objects, and it is specialized towards textural compositions of local shapes and colors — reminiscent of impressionist paintings. In contrast, BigGAN was trained to conditionally generate objects learned from ImageNet [11] with nuisance variations. BigGAN images often contain centered, object-like shapes in a photo-realistic style, while it struggles to create simpler images. Both generators create images with different underlying statistics (Fig. 1D). We compared the ability of neurons to drive the synthesis of optimal images, specifically measuring the success rate, firing rate activation, and the optimized images in the two spaces, in order to shed light on the coding principles behind ventral stream neuronal function.

**Fig. 1.**
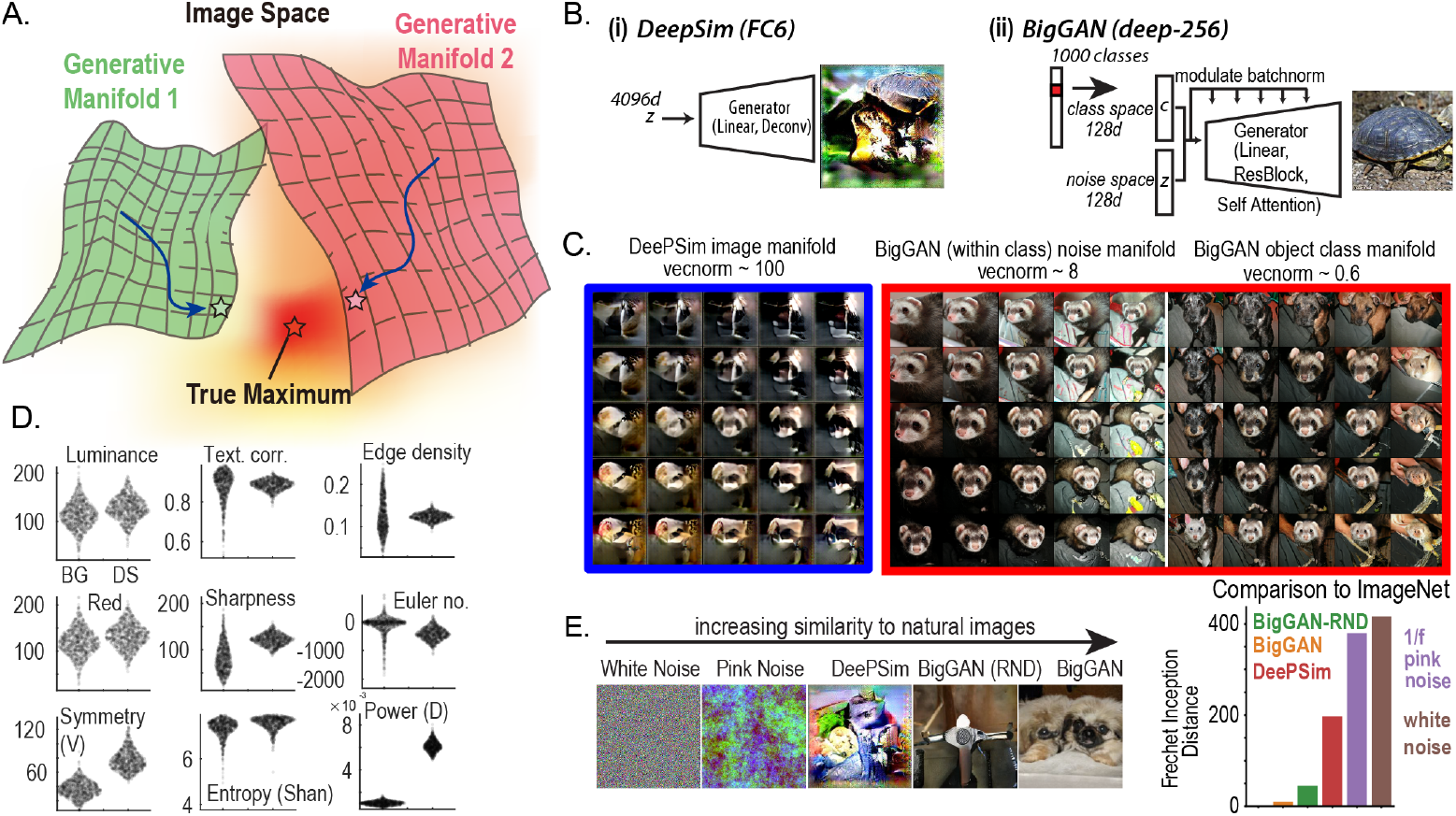
Image manifolds as parameterized by different generative networks (BigGAN and DeePSim generator). **A**. Conceptual schematic of complementary image manifolds as implemented in generative networks. **B**. Architectures of (i) DeePSim generator and (ii) BigGAN, highlighting the side modulation structure of BigGAN [12]. **C**. Image samples from each manifold: (left) **DeePSim-fc6**, (middle) **BigGAN**, samples from given class vector *c* while varying noise vector *z*; (right) **BigGAN**, samples from varying class vectors *c*. Each montage shares the same source image in the center. **D**. Low-level image statistics quantifying differences between DeePSim and BigGAN image samples. Each point represents a sampled image from the latent space of each generator (Tab. 5). **E**. Quantifying image similarity via *Frechet Inception Distance* between photographs (ImageNet validation images, N = 50,000) and other image manifolds, including *white noise* (i.i.d. uniform distributed pixel values), *pink noise*, following a 1/*f* spectrum, *DeePSim* images sampled from normally distributed latent vectors *z ∼ 𝒩* (0, 4*I*), and **BigGAN (RND)** sampled using normally distributed class vector *c* and noise distribution *z*, and **BigGAN**, sampled using trained class vectors and noise distribution values.

## 1. Results

### 1.1 Characteristics of the DeePSim and BigGAN image manifolds

To contextualize the biological results, we first characterized the properties of the image manifolds parametrized by the DeePSim and BigGAN generator.

DeePSim generator has a relatively simple architecture, comprising linear and upconvolution layers [7] (Fig. 1B). It has a single 4096-dimensional latent space, within which around 500 dimensions are enough to capture the generated image variations [8], with the rest effectively acting as a null space.

In comparison, BigGAN generator has a complex network structure, which includes self-attention [13] and modulated batch normalization[12]. It is also a class-conditional GAN (Fig. 1B), which receives a 128d class conditioning vector *c* and a 128d noise vector *z* . Each of the 1000 object classes in ImageNet is represented by a different class vector *c ∈* 𝒞; while the variations within each class are controlled by the noise vector *z ∈ Ƶ*, sampled from a spherical truncated Gaussian distribution during training. Thus, sampling closely related points (“traveling”) in the class embedding space 𝒞 interpolates between object categories, while traveling in the noise space *Ƶ* will change nuisance variables like aspect ratio, orientation, or viewing angle [10, 8] (Fig.1C, right). In comparison, traveling in the DeePSim space starting from the same image creates smooth pattern variations without preserving the central “object” (Fig.1C, left).

In general, the distribution of BigGAN images was closer to natural images than the DeePSim distribution was. We quantified this via the Frechet Inception Distance (FID) [14], a common metric of the distance between the image distributions of generative models (Fig. 1E).

We found BigGAN images sampled with 1000 pre-trained object vectors *c* were close to ImageNet images (FID 9.3). Since neuron-guided optimization could randomly change a given object vector, we also tested BigGAN images with Gaussian random object vectors (*BigGAN-RND*, FID 44.4), which were more naturalistic than DeePSim images (FID 197.0). In contrast, DeePSim images were less naturalistic, but still more so than pink noise (FID 379.5) and white noise (FID 416.2). Additionally, there were many differences in low-level image statistics between DeePSim and BigGAN images, as summarized in Fig. 1D and Table 5.

**Table 1.** Success rates of evolution experiments. Success criterion defined as an increase in activation from the first two generations (N = 25-40 images per generation) to two neighboring generations during the session, with statistical reliability estimated using a Student’s t-test, *P* < 0.01. Tables with alternative success criteria are shown in the Supplementary Information, Tab. 2, 3.

|  | BigGAN | DeePSim | Both |
| --- | --- | --- | --- |
| V1 | 0/10 (00.0%) | 10/10 (100.0%) | 0/10 (00.0%) |
| V4 | 20/38 (52.6%) | 37/38 (97.4%) | 20/38 (52.6%) |
| PIT | 65/106 (61.3%) | 67/106 (63.2%) | 52/106 (49.1%) |
| Total | 85/154 (55.2%) | 114/154 (74.0%) | 72/154 (46.8%) |

**Table 2.** Success rates of evolution experiments with alternative success criteria: t-test between the activations in the first two generations and two generations around the maximum activation, *init*2 < *max*2, *p* < 0.05.

|  | BigGAN | DeePSim | Both |
| --- | --- | --- | --- |
| V1 | 2/10 (20.0%) | 10/10 (100.0%) | 2/10 (20.0%) |
| V4 | 24/38 (63.2%) | 37/38 (97.4%) | 24/38 (63.2%) |
| PIT | 78/106 (73.6%) | 69/106 (65.1%) | 61/106 (57.5%) |
| Total | 104/154 (67.5%) | 116/154 (75.3%) | 87/154 (56.5%) |

**Table 3.** Success rate of evolution experiments with alternative Success criterion: t-test between the activations in the first two generations and last two generations *init*2 < *last*2, *p* < 0.01.

|  | BigGAN | DeePSim | Both |
| --- | --- | --- | --- |
| V1 | 0/10 (00.0%) | 10/10 (100.0%) | 0/10 (00.0%) |
| V4 | 15/38 (39.5%) | 35/38 (92.1%) | 14/38 (36.8%) |
| PIT | 54/106 (50.9%) | 59/106 (55.7%) | 41/106 (38.7%) |
| Total | 69/154 (44.8%) | 104/154 (67.5%) | 55/154 (35.7%) |

**Table 4.** Additional convergence time statistics for each visual area and statistical tests. **Upper**, mean±SEM of the statistics. Only threads with successful evolution in that space were included. (max > init, *p* < 0.01) **Lower**: Significant test for the comparisons presented in the main paper with alternative statistics. Independent t-test for IT vs V4 and IT vs V1, paired t-test for FC vs BG in IT.

| Area | V1 |  | V4 |  | IT |  |
| --- | --- | --- | --- | --- | --- | --- |
| GAN | FC (N=10) | BG | FC (N=37) | BG (N=20) | FC (N=67) | BG (N=65) |
| raw | 4.8±2.5 | | 7.9±1.2 | 6.0±1.7 | 12.7±1.1 | 7.4±1.1 |
| evoke | 7.0±2.4 | | 9.2±1.2 | 8.2±2.1 | 16.9±1.0 | 9.7±1.1 |
| bsl init | 10.1±3.1 | | 14.3±1.4 | 16.9±1.8 | 22.1±1.0 | 16.7±1.0 |

| Area | IT vs V4 | IT vs V1 | IT |
| --- | --- | --- | --- |
| GAN | FC (df=102) | FC (df=75) | FC vs BG (df=51) |
| raw | t=2.75, p=7.0e-03 | t=2.62, p=1.1e-02 | t=3.34, p=1.6e-03 |
| evoke | t=4.75, p=6.7e-06 | t=3.57, p=6.2e-04 | t=4.95, p=8.4e-06 |
| bsl init | t=4.67, p=9.3e-06 | t=4.40, p=3.6e-05 | t=3.36, p=1.5e-03 |

**Table 5.**
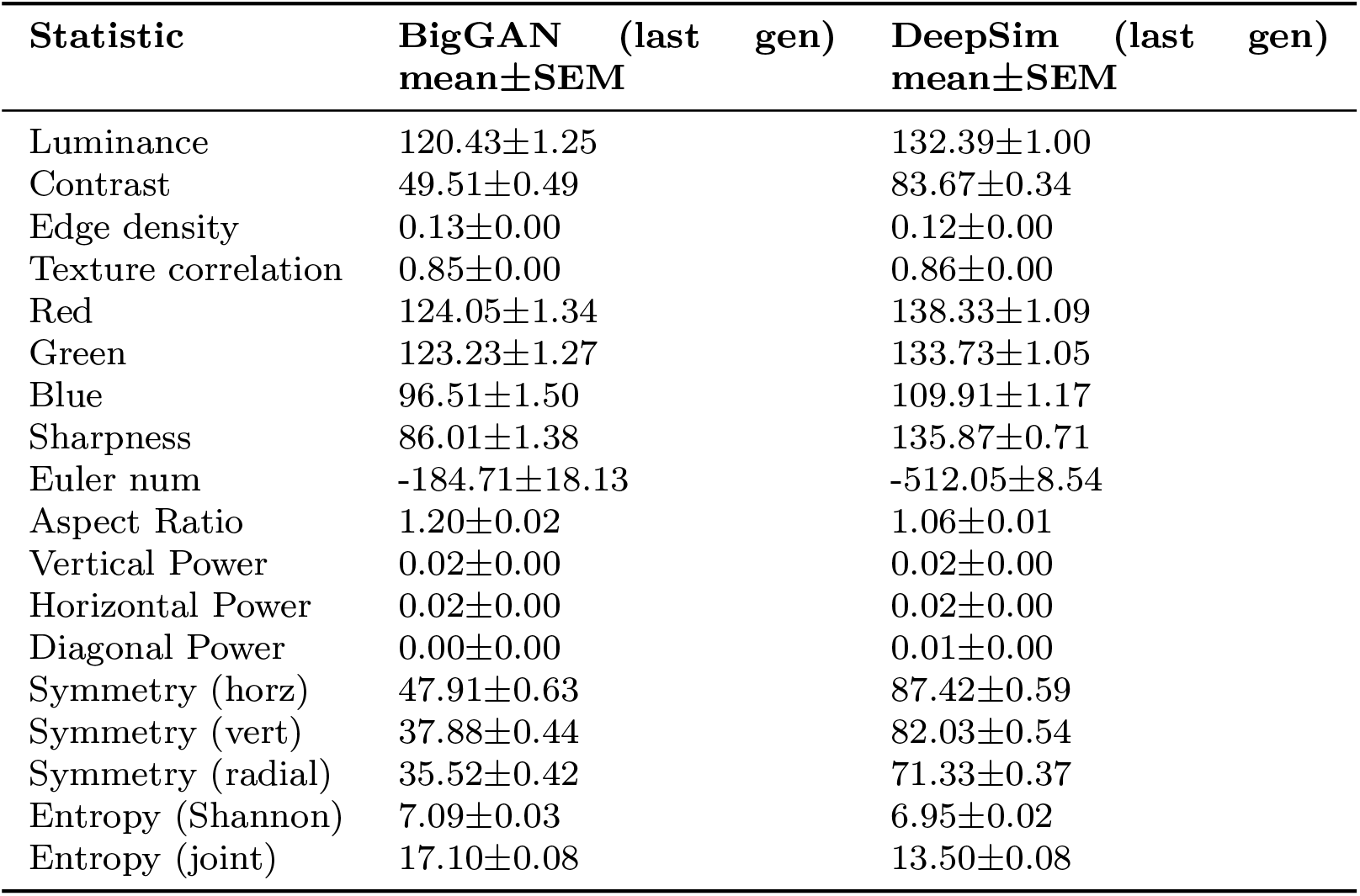
Low-level statistics of images optimized in DeePSim and BigGAN space, targeting the same hidden units in AlexNet.

| Statistic | BigGAN (last gen)<br>mean±SEM | DeepSim (last gen)<br>mean±SEM |
| --- | --- | --- |
| Luminance | 120.43±1.25 | 132.39±1.00 |
| Contrast | 49.51±0.49 | 83.67±0.34 |
| Edge density | 0.13±0.00 | 0.12±0.00 |
| Texture correlation | 0.85±0.00 | 0.86±0.00 |
| Red | 124.05±1.34 | 138.33±1.09 |
| Green | 123.23±1.27 | 133.73±1.05 |
| Blue | 96.51±1.50 | 109.91±1.17 |
| Sharpness | 86.01±1.38 | 135.87±0.71 |
| Euler num | -184.71±18.13 | -512.05±8.54 |
| Aspect Ratio | 1.20±0.02 | 1.06±0.01 |
| Vertical Power | 0.02±0.00 | 0.02±0.00 |
| Horizontal Power | 0.02±0.00 | 0.02±0.00 |
| Diagonal Power | 0.00±0.00 | 0.01±0.00 |
| Symmetry (horz) | 47.91±0.63 | 87.42±0.59 |
| Symmetry (vert) | 37.88±0.44 | 82.03±0.54 |
| Symmetry (radial) | 35.52±0.42 | 71.33±0.37 |
| Entropy (Shannon) | 7.09±0.03 | 6.95±0.02 |
| Entropy (joint) | 17.10±0.08 | 13.50±0.08 |

### 1.2 Adapting closed-loop image synthesis with BigGAN

Next, we set out to make sure that the closed-loop neural-guided image synthesis system [3, 6, 4] would work with newer generative networks such as BigGAN. This required developing appropriate *optimizers* to search in the latent space of the generator. Developing closed-loop optimizers for visual cortex neurons is costly, so we first simulated the process with a more tractable synthetic problem, a common strategy in the field of evolutionary computing [15, 9]. As image-computable models, convolutional neural networks (CNNs) serve as good approximations of the primate visual system[16]. We have shown that the tuning landscapes of CNN hidden units share many similar geometric properties as those of visual neurons [5]. Thus, we first tuned the configurations of various evolutionary algorithms for BigGAN via CNN hidden units. Because the process relies on evolutionary algorithms, we refer colloquially to the closed-loop optimization process as an *evolution*.

We identified a successful configuration of evolutionary algorithms that worked well for BigGAN *in silico*. The key parameter was the standard deviation of the sampling distribution (*i*.*e*., exploration step size). This parameter controls the distance of exploration in the latent space, which translates to the distances between image samples. If the images within a batch sample were too similar to each other, the difference in neuronal response to images would be overwhelmed by single-trial response variability, obscuring the “gradient” direction of the tuning landscape. On the other extreme, if the images within a batch were too far apart from each other, traveling in the averaged direction of their latent codes would not help. Optimally, the search step parameter should be tuned to suit the average *slope* of the neuron’s tuning function. Heuristically, we configured the search parameter for each generator such that the samples in each batch were perceptually different but showed some continuity. We tested these optimizers on *in silico* units from convolutional neural networks, and confirmed that they could successfully optimize activation for these units. We then tested these parameters *in vivo* (next section).

### 1.3 Neuron-guided image synthesis in two generative spaces

Having adapted this new generator for *in silico* optimization, we set out to compare how visual cortex neurons directed image optimization in each generative space. We conducted a series of parallel-evolution experiments using two male macaque monkeys (animals *A* and *B* ). The animals were implanted with chronic floating microelectrode arrays in three visual cortical areas (one at the border of V1/V2, immediately posterior to the lunate sulcus, another in V4, on the prelunate gyrus, anterior to the lunate sulcus, and the last in posterior IT, immediately anterior to the inferior occipital sulcus). Each day, after sorting and classifying the neuronal signals as arising from singleand multi-units (SU/MU), we mapped the approximate location of the receptive fields (RFs) of all recording channels and chose a visually responsive unit for study, which we call the *driver unit*. Using the response rate of the driver unit as the optimization target, we conducted two evolution experiments in parallel (*threads*), one searching within the BigGAN manifold, and the other within the DeePSim manifold, with separate optimizers configured for each space.

In each *block* or *generation*, the images proposed by the optimizer for BigGAN and DeePSim were randomly interleaved in a sequence, matching the recording quality and state of the animal between threads. Each image was shown once to the monkey for 100-ms *on*, 150-ms *off*, 3-5 images per trial. The monkey was required to maintain fixation in a circle of 1° radius. Failure to maintain fixation aborted the trial. After all images were presented, the mean firing rate of the driver unit in 50-200ms to each image was summarized as a scalar score and provided to the optimizers (see **Methods**). Then, the optimizers proposed two new sets of latent vectors, which were rendered into images, initiating the next block (Fig. 2A).

**Fig. 2.**
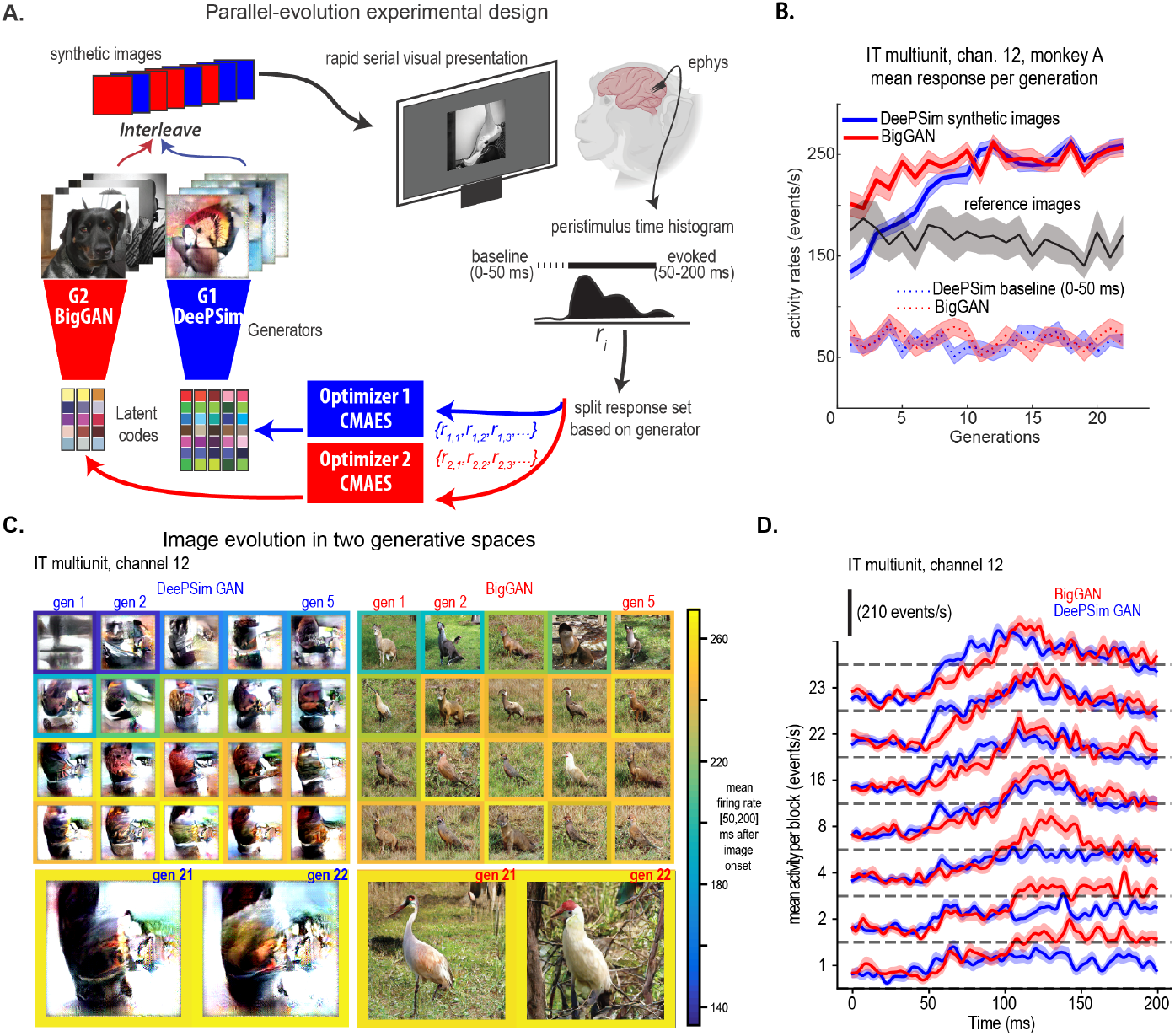
Example of a successful paired evolution from an inferotemporal cortex site. **A**. Schematic showing the workflow of a parallel evolution experiment *in vivo*. **B**. Neural activation during paired evolution in an example session for PIT multi-unit. Line and shading show the mean firing rate and standard error (SEM) . **Colored solid lines** show the evoked firing rate per block, as generated via the BigGAN (red) and DeePSim (blue) spaces. The **gray solid line** shows the firing rate for a fixed set of (non-evolving) reference images. The **colored dotted lines** show the baseline firing rate (0-50ms) for each thread. **C**. Evolved images per the DeePSim and BigGAN spaces. Example images evoking the highest firing rate in each block are shown. Color frames illustrate the average evoked firing rate of the images per block. **D**. Evolution of response dynamics per the peristimuli time histogram, (PSTH) across generations. Lines and shading show the mean and SEM of PSTHs across images in each block.

We collected 170 paired evolutions, but excluded experiments where the recording quality was not stable (*e*.*g*., with fluctuating baseline firing rates, as measured via non-evolving reference images) or if the experiment had fewer than 15 blocks. This resulted in a dataset of 154 sessions (90 in monkey A, 64 in B).

We start by focusing on a typical experiment involving a PIT multi-unit site in monkey A. Both DeePSim-and BigGAN-based algorithms successfully increased the neuronal firing rate (Fig. 2B,C). The corresponding optimized images showed different global configurations, but shared local motifs. For DeePSim, the image optimization started from a typical texture, and the preferred feature was “synthesized” from scratch. In contrast, for BigGAN, the image optimization started by choosing a well-formed alpaca-like creature on a green background, and then the object identity and nuisance variables were gradually refined to maximize the response. After response convergence, in DeePSim space, the neuron guided the generation of a brown curved surface against a white background; while in BigGAN space, the neuron guided the generation of a bird-like creature on a green grassland background (Fig. 2D.). Below, we will show that in general, neurons guided both generators to synthesize local features, such as the curved edge present in the bird’s neck, which is well-isolated in the optimized DeePSim images. Although these optimized images were globally different, they evoked comparable firing rates: (mean firing rate in [50, 200]-ms window for images of the last two blocks; two-sample t-test *t*_128_ = 0.97, *P* = 0.33). This suggests that the neuron could respond similarly to two globally different images as long as key features were embedded in both of them. Inspecting the mean peristimulus time histograms (PSTHs), we found interesting differences in the neural dynamics evoked through the BigGAN and DeePSim spaces. For example, the response peaks for the BigGAN images showed a longer latency than those for the DeePSim images (Fig. 2 C). We will analyze this effect below.

### 1.4 Success rates highlight a differential alignment between visual area and generative spaces

Conceptually, we might regard the facility with which neurons guide the optimization as the *alignment* between the neuronal tuning and the latent space of the generative model. If the neurons were smoothly tuned to latent variables controlling the images, a more aligned tuning might create a smoother tuning landscape, facilitating hill climbing. To investigate this alignment, we analyzed the facility for neurons in each visual area to guide the evolution process on each image manifold, and we quantified it through the success rate across evolution experiments. We defined an evolution as successful if the neuronal activity rate was statistically higher in two neighboring blocks than it was at the first two blocks (mean firing rate in 50-200 ms for 25-40 images per block), as statistically measured using a Student’s t-test (Tab.1). Using this criterion (*p <* 0.01), we found that across all areas, the overall success rate of BigGAN evolutions (55.2%, 5-95% confidence interval [48.6%, 61.6%]) was lower than that of DeePSim evolutions (74.0%, CI [67.8%, 79.3]). Under a less strict statistical criterion (*p <* 0.05), the overall success rates were comparable: DeePSim, 75.3%, CI [69.1%, 80.5%] vs BigGAN, 67.5%, CI [61.0%, 73.3%]. (Tab. 2)

When measuring the success rates conditioned on the cortical area of the driver unit, we found that the BigGAN and DeePSim threads showed different patterns (Tab.1). For BigGAN evolutions, the success rate increased from V1 (0*/*10), through V4 (20*/*38), and to PIT 61% (65*/*106). In contrast, for DeePSim evolutions, the success rate fell from V1 (10/10, 100%) through V4 (37*/*38), to PIT (67*/*106, Fig. 3A). We replicated this analysis using other metrics for success, and while the overall rates could change, the trends across areas remained the same (Tab.3)). So, there was a clear gap between the success rates of DeePSim and BigGAN in both V1 and V4, while their success rates were comparable in PIT. Further, for PIT experiments, the evolution successes across both spaces were not independent: evolution success in one space was statistically associated with success in the other space (per a Chi-square contingency test, 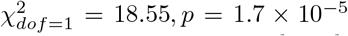, Fig. 3B), suggesting that a common property such as responsiveness or signal quality partially determined the optimization success in both spaces. Overall, these results showed that visual cortex neurons could guide different types of image generators to synthesize highly activating stimuli, consistent with the views that visual cortex neurons have smooth, continuous tuning on both image manifolds that allowed their respective optimizers to “climb” slopes in each space. The differences in success rates across the visual hierarchy suggested that the pattern-based parametrization of DeePSim was more aligned with the neuronal tuning in V1 and V4, while both the object-based, photo-realistic parametrization of BigGAN was more aligned to a deeper stage in the hierarchy.

**Fig. 3.**
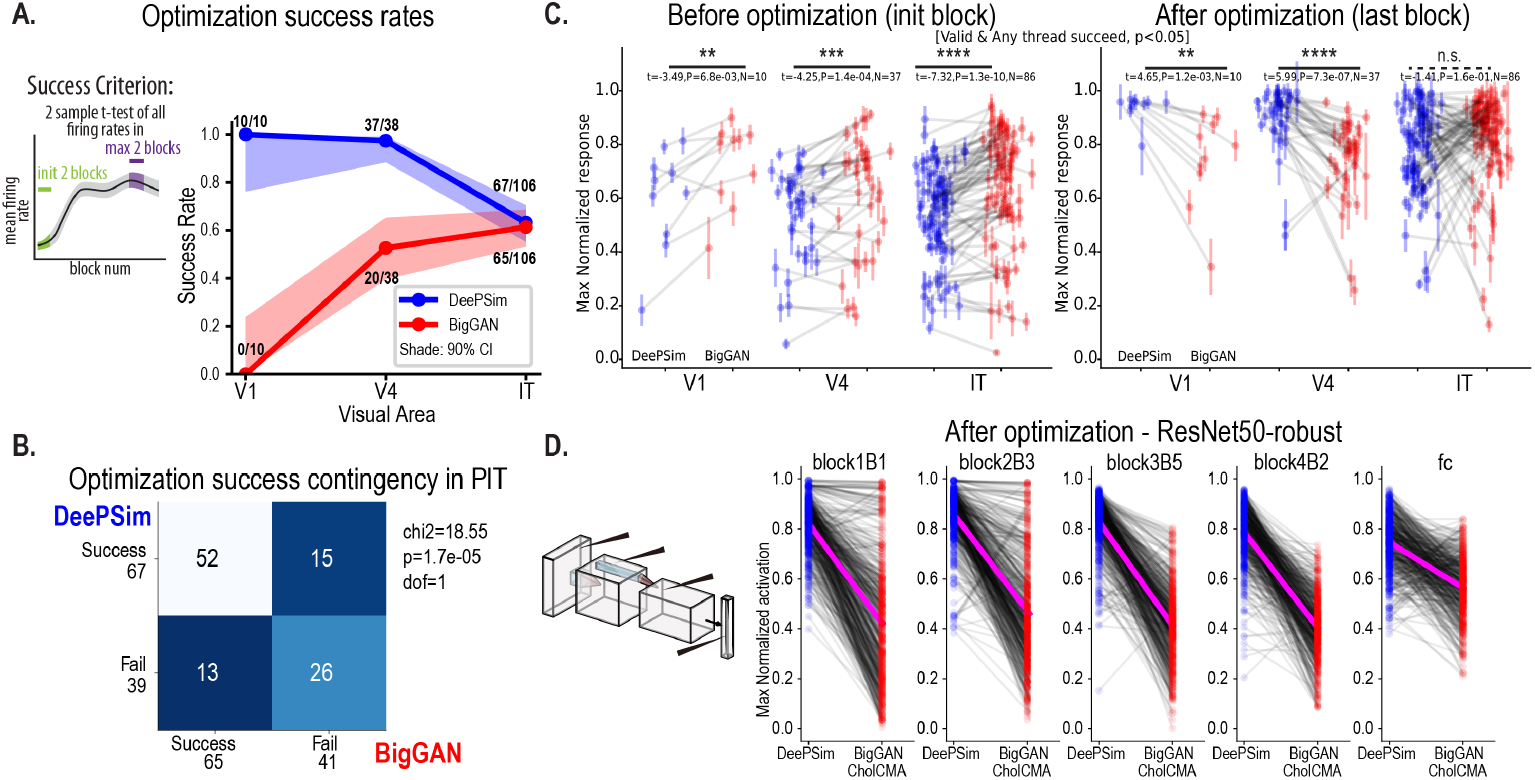
Evolution success rates and neuronal activity levels across DeePSim and BigGAN spaces. **A**. Evolution success rates for sites in V1, V4, and PIT (*p* < 0.01 criterion, max > initial activation). Left, schematics of the success criterion. **B**. Contingency table showing evolution successes per image space are correlated. **C**. Neuronal activity rates before (left) and after optimization (right). The dots and error bars show the mean and SEM of neural activation in the initial or final block, with paired threads connected. Activity rates were normalized by the maximum block-wise mean activation across blocks and threads, experiments where any thread succeeded were included (Supplementary Figure 8 shows outcomes when only experiments where both threads succeeded were included). **D**. Maximum normalized activation after optimization (last block) in DeePSim and BigGAN space for *in silico* experiments on ResNet50-robust. Parallel evolution experiments were performed on 50 units in given layers across the hierarchy with 10 repetitions each.

### 1.5 Visual areas showed different maximal activations depending on image space

To explore how visual areas aligned to different image spaces, we analyzed the activity evoked by BigGAN- and DeePSim images before and after neuron-guided optimization. We found that before optimization (in the first generation), BigGAN images generally evoked a higher firing rate across all three visual areas (BigGAN > DeePSim, paired t-test, max > init *p* < 0.05, *t*_9_ = −3.491, *p* = 6.8 × 10^−3^ for V1, *t*_37_ = − 4.399, *p* = 8.9 × 10^−5^ for V4, *t*_105_ = − 9.286, *p* = 2.4 × 10^−15^ for PIT, N = 154 experiments). This suggests that the default visual statistics learned by BigGAN are well-suited to visual neurons, in contrast to the initial textures learned by DeePSim (Fig.3C). However, after optimization, the results differed among visual areas. For experiments where at least one thread succeeded, DeePSim images evoked higher activations for V1 and V4 neurons (*t*_9_ = 4.651, *p* = 1.2 × 10^−3^ for V1, and *t*_36_ = 5.985, *p* = 7.3 × 10^−7^ for V4); for PIT neurons, BigGAN images generally evoked a similar level of activation as DeePSim(*t*_85_ = − 1.41, *p* = 0.16, *N*.*S*.; in monkey A, the optimized BigGAN images evoked a slightly higher activation for IT neurons (*t*_47_ = − 2.32, *p* = 0.025), and in monkey B, they were comparable *t*_37_ = 0.70, *p* = 0.49) (Fig.3C). One confounding factor is that the success rate differed for BigGAN and DeePSim. To control for this, we considered the Evolution experiments where both threads succeeded (N=87). In this case, for V4 neurons, the optimized activation for DeePSim space remained higher than BigGAN (DeePSim mean *±* sem 0.898 *±* 0.028, BigGAN 0.784 *±* 0.020, *t*_23_ = 3.707, *p* = 1.2 × 10^−3^); for PIT neurons, the optimized activations of two spaces were still comparable (DeePSim, 0.821 *±* 0.018, BigGAN 0.820 *±* 0.021, *t*_60_ = 0.06, *p* = 0.95) (Supplementary Fig.8). So, similar to the results above pertaining to success rate, the optimization gap between DeePSim and BigGAN was closed by PIT cortex.

Next, we conducted similar parallel evolution experiments on convolutional neural networks (CNNs). Specifically, for each CNN model, we selected driver units with receptive fields (RFs) at the center of the feature map and applied the same paired evolution experimental design as above. We ran 10 “sessions” per generative model (see *Methods*). We found that in contrast to visual cortex, units throughout the CNN hierarchies were almost always more activated by DeePSim-optimized images than by BigGAN-optimized images (Fig. 3D, DeePSim > BigGAN, paired t-test, *t*_499_ > 24.7, *p* < 4.7 × 10^−89^ for all layers). Yet, as in visual cortex, we found that this activation gap depended on the depth of the unit in the hierarchy: units in deeper layers exhibited a smaller activation gap. For example, in ResNet50-robust, units in the final object classification layer (fc) and the penultimate layer (block4B2) showed smaller gaps between the DeePSim/BigGAN activations than earlier layers (Fig. 3D, two-sample t-test, DeePSim - BigGAN gap in block1,2,3,4 > gap in fc, *t*_998_ > 16, *p* < 5 × 10^−55^, for all earlier blocks).

In summary, across areas, neurons showed different levels of optimized activity depending on image space. For V1 and V4 neurons, optimization in DeePSim space led to a higher activation than in BigGAN space, despite that it started from a lower activation. We interpret this as evidence that the tuning of V1 and V4 neurons is more aligned with a local-pattern-based image parametrization, where the tuning landscape peaks of these neurons project closer to the DeePSim manifold than to the BigGAN manifold. For PIT neurons, the optimization in the two spaces converged to comparable values, showing an increasing alignment with both DeePSim and BigGAN manifolds. This alignment was stronger in PIT neurons than it was in CNNs, even at the classification layer, highlighting a discrepancy between the visual stream and its most promising computational models.

### 1.6 Optimization dynamics depended on image space

We asked how neurons changed their activation over the evolution process, to estimate their *optimization trajectory*. Intuitively, neurons might exhibit different tuning landscape geometries on each manifold, which would affect the optimization dynamics. We hypothesized that if a given manifold contained the basic motifs encoded by a visual neuron — if the space *aligned* better with it — then the evolution should converge faster within that space. We computed the average optimization trajectory for each visual area (Fig. 4A). As shown above, for V1 and V4 neurons, even though BigGAN image space started with a higher firing rate, it was rapidly surpassed by the DeePSim space. In contrast, for PIT neurons, BigGAN space on average stayed on top and reached similar level of activation with DeePSim in the end.

**Fig. 4.**
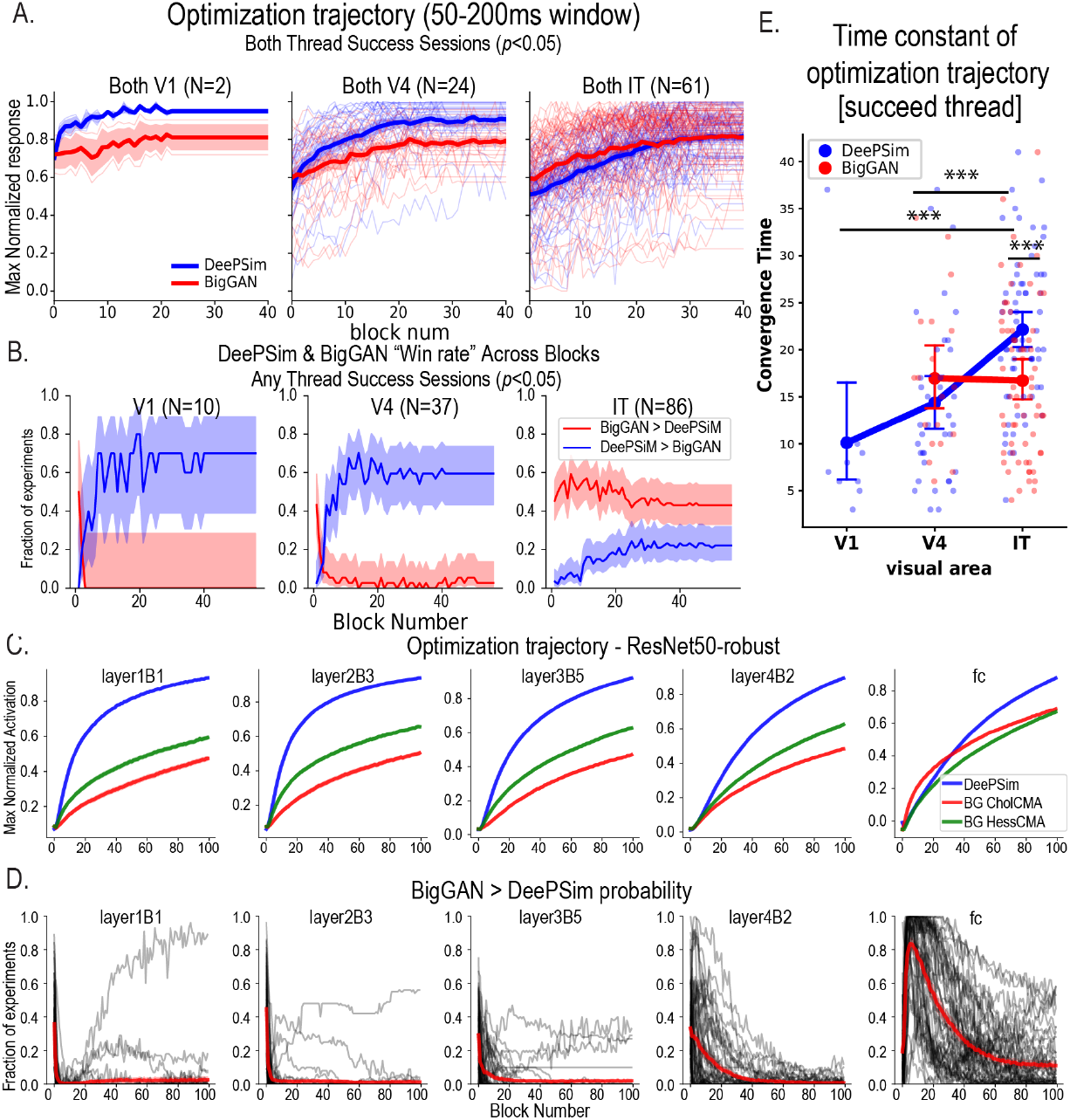
Optimization dynamics modulated by visual area and image space. **A**. Maximum normalized optimization trajectory for each image space averaged per visual area in sessions where *both* threads succeeded. Here we show results with a more inclusive success criterion with *p <* 0.05, since otherwise no session would be included for V1. **B**. Win rate of DeePSim and BigGAN as a function of blocks for V1, V4 and IT neurons. Sessions where any thread succeeded were included. Lines and shading show the win fraction and its 95% confidence interval, estimated through the Beta distribution. **C**. Average activity during DeePSim and BigGAN evolutions for units in ResNet50-robust; each unit’s activation was normalized by its maximum activation. The shaded error bar showing the SEM was covered by the line. **D**. Win rate of BigGAN for ResNet50-robust units. The thin black trace is the averaged win rate for each unit across 10 repetitions, the red curve is the average win rate curve for each layer averaged across all the units. **E**. Evolution convergence time constant for both spaces across visual areas, including all threads that succeeded. Small dots show the time constant of each successful thread, big dots and error bars show the mean and SEM of each area per space.

We quantified this through the *win rate* of each space across optimization. The win rate was defined as the fraction of sessions where neuronal responses to BigGAN images were significantly higher than those to DeePSim images (per Student’s t-test) or vice versa (Fig. 4B), for a given generation. For V1 and V4 neurons, initially BigGAN images evoked higher activation in around 40% sessions; but only after 2-3 blocks the BigGAN win rate dropped to around 0, and 60% of DeePSim threads climbed to higher activity. In contrast, for PIT neurons, throughout evolution, more than 40% of experiments BigGAN thread attained higher activation than DeePSim.

We repeated the same analyses for CNN units, and also found that for deeper units in the CNN, while the DeePSim evolution eventually reached higher activations, it took more blocks for the mean DeePSim-evoked activation to surpass the mean BigGAN-evoked activation (Fig. 4C,D. See also Supplementary Fig. 9).

In previous work, we found that neurons deeper in the visual hierarchy took longer to converge [4, 5], suggesting a sharper and higher-dimensional tuning geometry. Here, we quantified how long it took for evolution to reach convergence for both spaces.

Specifically, we measured the *time constant* of each successful evolution, namely the number of generations required to reach 80% of the maximum activation increase from the initial generation (Fig. 4E). In DeePSim space, the number of generations required to find optimized images increased along the visual hierarchy (PIT > V4 22.1 ± 1.0 (*N* = 67) versus 14.3 ± 1.4 (*N* = 37), *t*_102_ = − 4.67, *p* = 9.3 × 10^−6^, PIT > V1 10.1 ± 3.1(*N* = 10), *t*_75_ = − 4.40, *p* = 3.6 × 10^−5^), consistent with our previous findings. Further, when both threads succeeded, for PIT neurons, optimization in the BigGAN space converged faster than in the DeePSim space (DeePSim > BigGAN 22.3 ± 1.1 vs 17.7 ± 1.1 (*N* = 52), *t*_51_ = 3.36, *p* = 1.5 × 10^−3^)). Results with alternative convergence measures are presented in Tab. 4.

Overall, these results suggest a stronger alignment of PIT neuronal encoding to the BigGAN manifold compared to V1- and V4 neuronal encoding. The continuing effectiveness of the DeePSim manifold in PIT also suggests that the BigGAN manifold is not uniquely preferred by these higher visual neurons.

### 1.7 BigGAN aligned best to late neuronal responses in PIT

We have shown that biological neurons in PIT cortex showed stronger alignment towards the object-based generator (BigGAN) compared to CNN units. To understand this, we explored one feature that biological neurons have but most classic discriminative models of vision lack: temporal dynamics. Visual neurons show dynamic responses to static images. These dynamics have been known to reflect shifts in the encoding of visual features [17,18,19,20,21,22]. Dynamics may reflect feature selectivity changing from broader to sharper tuning, from spatially local tuning to holistic or contextual, or from coarse to fine-grained information. Here, we analyzed an elementary form of response dynamics, *i*.*e*., the peristimulus time histogram (PSTH) throughout each evolution (Fig. 2D). We analyzed (1) the response after image onset (PSTH, response as a function of time in milliseconds), and also (2) the time-averaged response over the image optimization session (response as a function of blocks during the evolution).

How did the dynamics of neuronal tuning interact with the optimization objective? By design, the optimization objective of our experiment was to maximize the overall firing rate in the [50, 200]-ms time window of the PSTH, but it was the “choice” of the neuron to determine *how* this would occur: the algorithm might increase the early, transient response, the late sustained response, an off response, or all of these stages together. To determine what changed, we examined the overall activation increase in different time windows after image onset. For all successful evolutions (criterion of *p* < 0.01, two-sample t-test), we computed the firing rate difference in each 10-ms time bin as a fraction of the total increase of mean firing rate (0-200ms) (Fig. 3G). We found that in PIT, the activation increase in BigGAN could be attributed more to later periods, [100, 110]-ms (DeePSim > BigGAN, two-sample t-test, same below, *t*_130_ = − 2.129, *p* = 0.035) and [120, 130] ms (*t*_130_ = − 2.967, *p* = 0.0036) and less to initial periods, [50, 60] ms (*t*_130_ = 2.144, *p* = 0.034) and [60, 70] ms (*t*_130_ = 2.574, *p* = 0.011). This showed that successful BigGAN evolutions preferably recruited a later period of neuronal response for PIT neurons.

Further, how did the PSTH dynamics interact with optimization dynamics in the session? We computed the firing rate in 25-ms time bins after image onset, and tracked activation in each bin throughout the session blocks, which we called a *time-split optimization trajectory*. We found that, for V4 neurons, the response to the DeePSim image surpassed that of the BigGAN images in all the 25-ms time windows from 50ms to 150ms (Fig. 5F, upper) and comparable for [150, 200]ms. However, for PIT neurons, in the initial time windows ([50, 75]-ms, [75, 100]-ms), the optimization in DeePSim space increased the firing rate more than that in the BigGAN space; but in later time windows (starting 100ms) BigGAN images started higher and stayed over DeePSim images (Fig. 5E, lower).

**Fig. 5.**
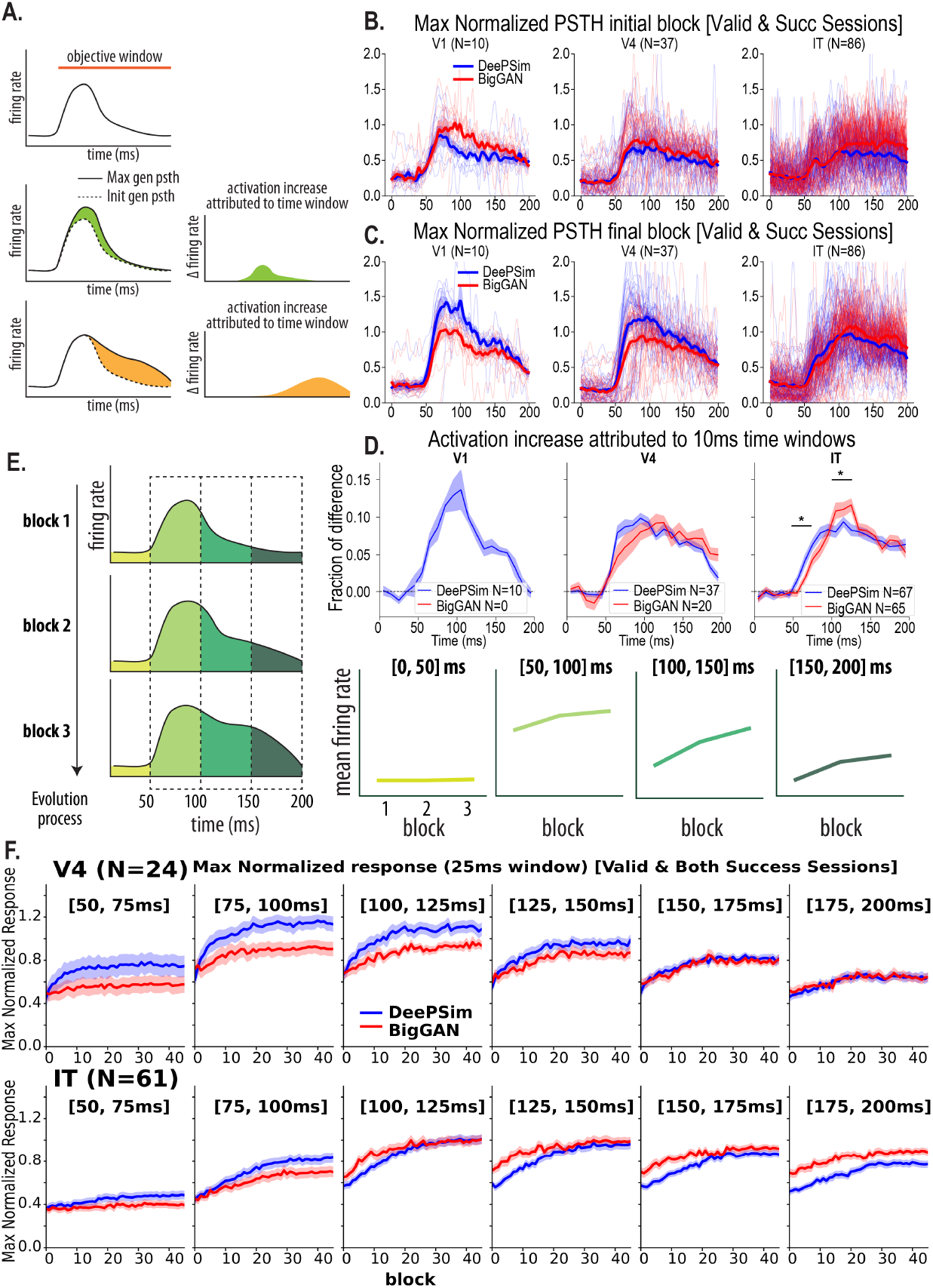
BigGAN space preferably activated and recruited late response in PIT neurons.. **A**. Schematics showing the analyses of neural dynamics in evolution experiment: upper, tracking activation increase in each time bin through evolution; lower, attribute activation increase to different bins. **B.C**. Averaged PSTH of units in V1, V4 and PIT before (**B**.) and after (**C**.) optimization. The thin lines showe the block-averaged PSTH for one thread, the thick lines and shading show the mean and SEM of PSTH across threads. **D**. Activation increase attributed to 10-ms time bins, successful threads in each space were included. Lines and shading show the mean and SEM across successful threads for V1, V4 and IT neurons. **E**. Schematics of the time-window specific Evolution trajectory analysis: evoked firing rates were computed for fine-grained time windows, then these firing rates were tracked as a function of blocks as evolution trajectory. **F**. The evolution trajectory of V4 and PIT neurons with firing rates in different 25ms time windows, mean and SEM of trajectory across neurons are shown. For PIT neurons, in the earlier time windows, the DeePSim evolutions prevailed and surpassed BigGAN throughout the blocks; however, in the later time windows, BigGAN surpassed DeePSim throughout the Evolution.

In summary, even though the mean activity after optimization was comparable in DeePSim and BigGAN space for PIT neurons (Fig. 3C), the dynamic responses differed across spaces. For PIT neurons, BigGAN images evoked stronger activity in later stages(Fig. 5F); also, more late responses were recruited while optimizing BigGAN images compared to DeePSim images (Fig. 5D). Both of these results suggested a better alignment of the late response tuning of PIT neurons with the object-parameterized image manifold.

### 1.8 Image similarity was related to neuronal dynamics

Next, we asked if a given unit drove the synthesis of similar visual features in both generative spaces. We note that there is no single metric of similarity: an individual’s perceptual system defines one measure of similarity, each given computational vision system defines another. Importantly, each neuron or neuronal microcluster also defines its own objective metric of similarity. As such, if neurons showed comparable levels of activation to different images in each generative space, then by definition, those images would be similar to the neuron. Here, we wanted to identify pixel-level features that could be related to that neuronal similarity. Visual examination of the optimized images from each generative space suggested they were globally different but shared similar local motifs. For example, one PIT microcluster (monkey A) used the DeePSim generative space to drive the synthesis of an upward-curving contour, segmenting a brown shape against a white background. The same site used the BigGAN space to drive the synthesis of an egret-like bird with a curly neck against the lighter yellow-greenish background, resembling the same contour (Fig. 2D). Generally, the experiments revealed the explicit biases of the generative networks: DeePSim was powerfully expressive, generating abstract patterns that often lacked globally cohesive shapes, whereas BigGAN was considerably less abstract due to its bias towards creating sharply focused objects, even for V1 (Fig. 6A). However, the neuronal responses could be quite similar for said images (Fig. 6B). To quantify image similarity, we performed two analyses: one using the feature space of a pre-trained image encoder (*e*.*g*., ResNet50-robust [23, 24]) to obtain an overall similarity score, and the other analysis using a network based on perceptual similarity (LPIPS, [25]) to obtain the spatial similarity maps of local patches (Sec. 1.9).

**Fig. 6.**
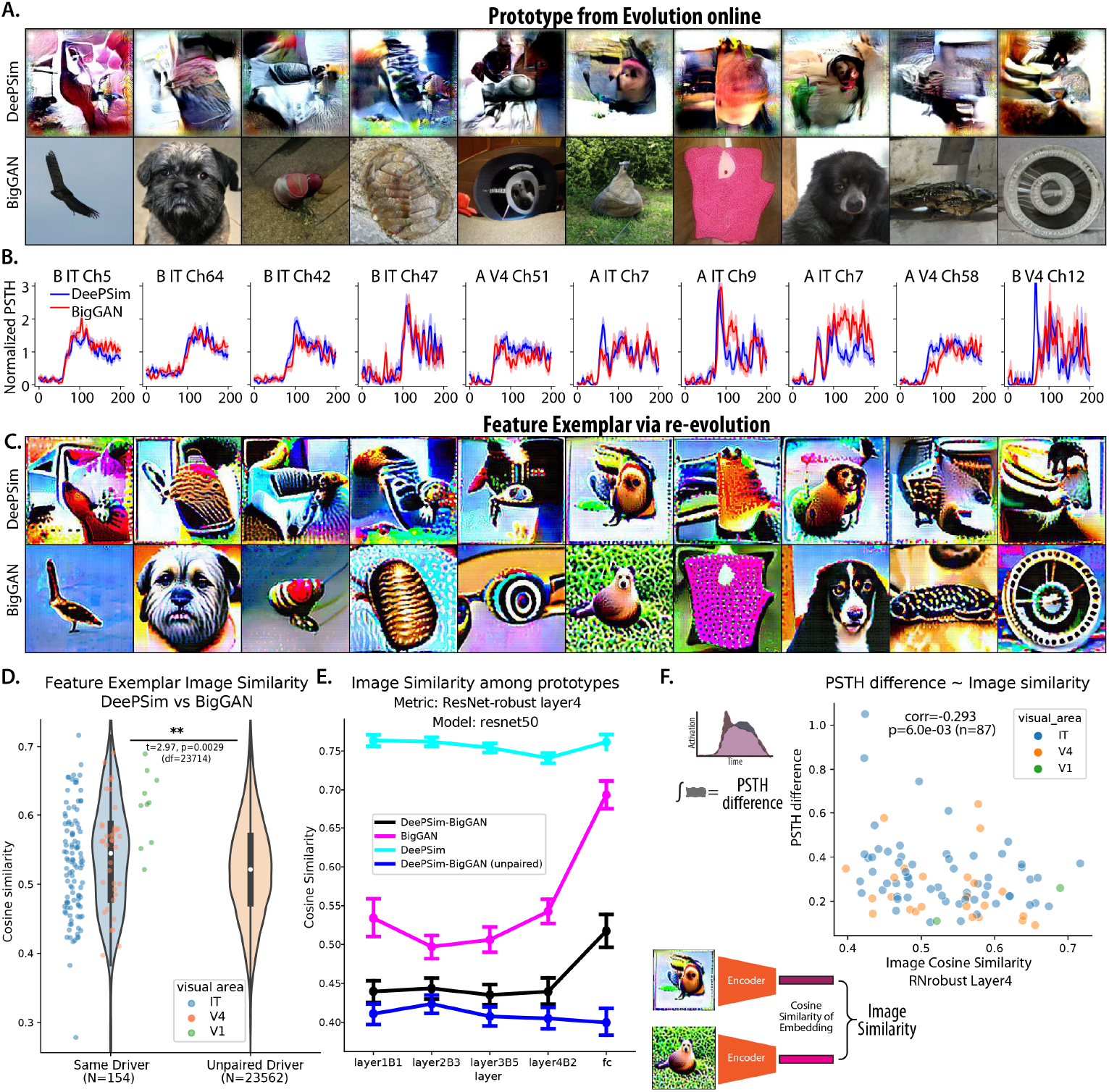
Similarity of evolved features in the two generative spaces, conditioned on evolution success. **A**. Selected evolved images from DeePSim and BigGAN. Images that evoked the highest activations were selected. **B**. Comparison of PSTH for the image block evoking the highest firing rate in DeePSim and BigGAN evolution, lines and shading show the mean and SEM of PSTH in that block. **C**. Feature exemplars obtained by re-optimizing the computational model of the recorded evolution. **D**. Image similarity of paired DeePSim and BigGAN prototypes from *in vivo* evolutions was higher than the similarity of shuffled pairs. The inner box plots show the median and quartiles of the distribution. The scatter plots show the distribution of prototype pairs for each area. All sessions were included. For other similarity metrics, see Supplementary Figure 10. **E**. As in **D**, but applied to evolution of *in silico* units from ResNet50. The error bar shows the 95% confidence interval of the mean, across pairs. For other computational models, see also Supplementary Figure 11. **F**. The similarity of the optimized image was anti-correlated with the distance of the neural PSTH.

For the first analysis, we first tried to identify the most important features contained within the evolved images, using a model-based approach. Evolution experiments resulted in a sequence of images and their evoked responses. For each evolution thread, we used all images and neuronal responses to fit a predictive model of the neuron, and then optimized a new image *in silico* in pixel or DeePSim generator space to drive this model unit. We called the resulting image from *in silico* optimization the *feature exemplar*, and frequently, this re-evolution emphasized key features more saliently than the original single images [26] (see Methods, Fig. 6C). With these images, we then explored if the images synthesized from dual-evolution were generally similar to each other, by passing them through a pre-trained encoder and computing the feature similarity in the embedding space.

We found that generally, the similarity between the maximal activating images in DeePSim and BigGAN space for the same driving unit was higher than that for unmatched drivers (ResNet50-robust layer 4 embedding, paired vs unpaired, mean ± std 0.414 ± 0.084 (N=154) vs 0.397 ± 0.078 (N=23562), independent t-test *t*_23714_ = 2.61, *p* = 9.1 × 10^−3^). When this measure was applied to the re-evolved feature exemplar, the similarity was higher and the effect size more prominent (paired vs unpaired, 0.539 ±0.076 (N=154) vs 0.521 ± 0.073 (N=23562) *t*_23714_ = 2.97, *p* = 2.9 × 10^3^, Fig. 6D). Image similarity metrics using other pre-trained encoders (e.g. AlexNet, VGG) yielded qualitatively the same result (Supplementary Fig. 10). The same analyses were conducted on the evolution driven by units *in silico* (Fig. 6E). Similar to neurons, the prototypes generated in two spaces by the same hidden-unit driver were more similar than pairs with unmatched drivers. More generally, when driven by the same units, the prototypes generated in one space were more similar to each other than pairs between spaces, with prototypes within DeePSim space more consistent.

Next, we focused on the dynamics of this emerging similarity. We computed the mean PSTH during the maximum-activation block in each generative space and compared these paired PSTHs. We found that the integrated absolute difference between the two normalized PSTH (*d*_PSTH_) was a good predictor of the image similarity between both spaces (*c*_embed_). For paired evolutions where both threads succeeded, the Pearson correlation between *d*_PSTH_ and *c*_embed_ was − 0.293, (*N* = 87, *p* = 0.006) (Fig. 6F). We found that the PSTH distance was a better predictor of image dissimilarity than the difference between scalar activation *d*_act_, which had a Pearson correlation of *c*_embed_ − 0.098, (*N* = 87, *p* = 0.37, *N*.*S*.). This showed that the specific time-course differences between PSTH encoded important information about the difference between the two prototypes.

### 1.9 Image similarity was local

Finally, we asked if the similarity between images was local vs. configurational. Per visual examination, it appeared as if neuronal sites led to globally different images with comparable local regions. For example, in our first experiment, IT site #20 in monkey B drove the evolution of two images from very different regions in image space (Fig. 7A,B): the final BigGAN image resembled a black side-view car mirror against a blue background, and the DeePSim image looked like a spread-out colored texture. Both images were equally activating (Fig. 7C), and both threads optimized images shared a centrally placed, black smooth oval against a pink smooth background, frequently present in monkey faces and animal ears (Fig. 7B). To test if these regions were similar, we used the LPIPS library to create a similarity map between the images (similarity = 1 – LPIPS distance). We saw that the map was not homogeneous but acquired high values in local regions. To determine if these local regions were likely to arise by chance across unrelated images, we measured the similarity maps between BigGAN – DeePSim image pairs optimized for the same unit (same-driver maps, Fig. 7D, top row) and compared them against the heatmaps between BigGAN–DeePSim image pairs optimized for different units (different-driver maps, Fig. 7D, bottom row).

**Fig. 7.**
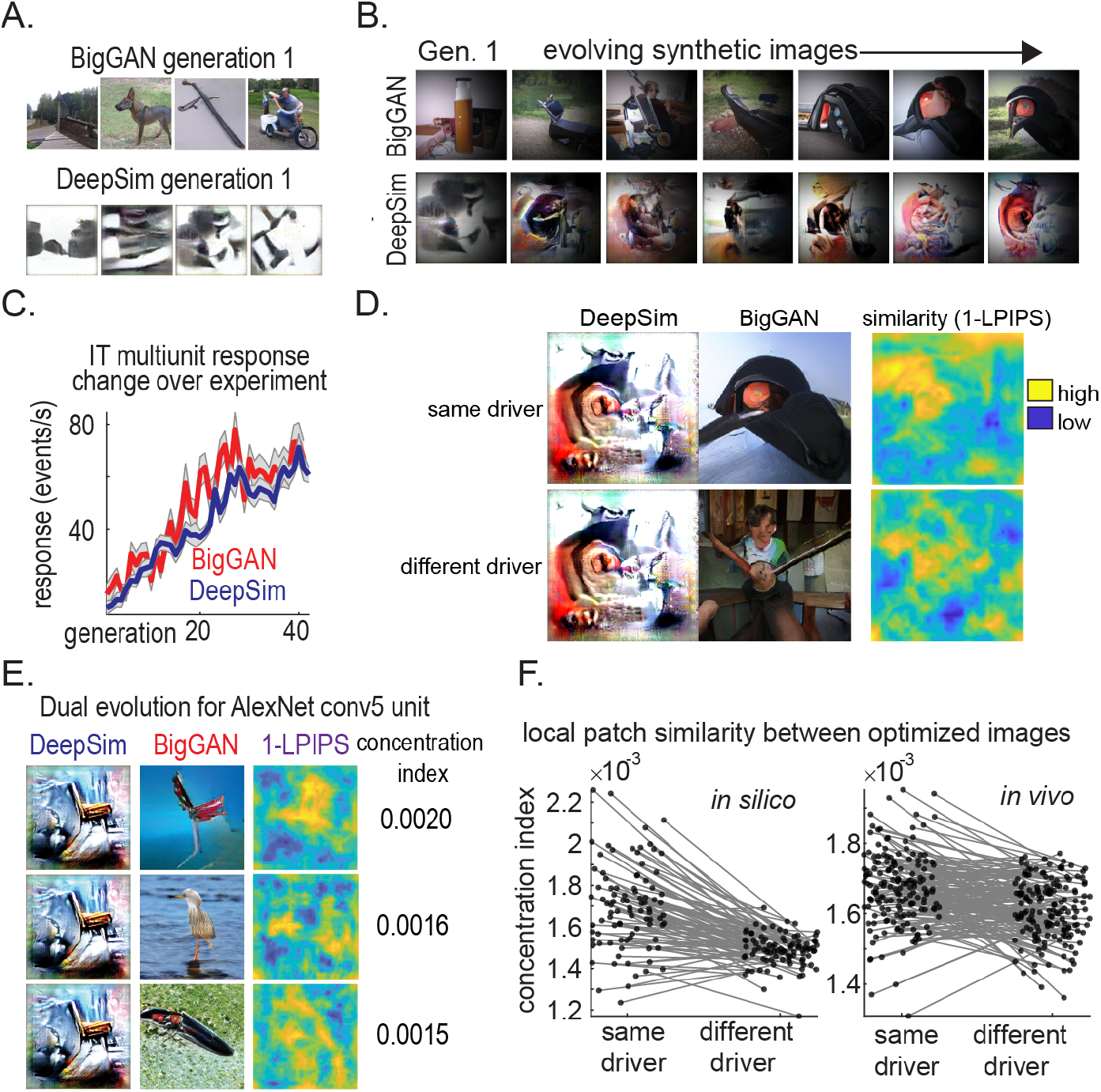
Images were locally similar. **A**. Sample of generation-0 images used for the BigGAN and DeePSim evolutions. **B**. Sample images showing changes over the experiment. **C**. Neuronal microcluster activity (mean firing rate per generation, ± S.E.M) in DeePSim (blue) and BigGAN (red) evolutions. **D**. LPIPS similarity map between optimized images for the same PIT site (top row) vs. optimized images for different sites (bottom row). **E**. LPIPS similarity map between optimized images for the same *in silico* units (AlexNet *conv5* )vs. optimized images for different units, along with an index showing local similarity concentration. **F**. Concentration index of similarity among BigGAN–DeePSim images when driven by the same unit (“same driver”) and when driven by different units (“different driver”), for both CNN- and visual cortex sites.

**Fig. 8.**
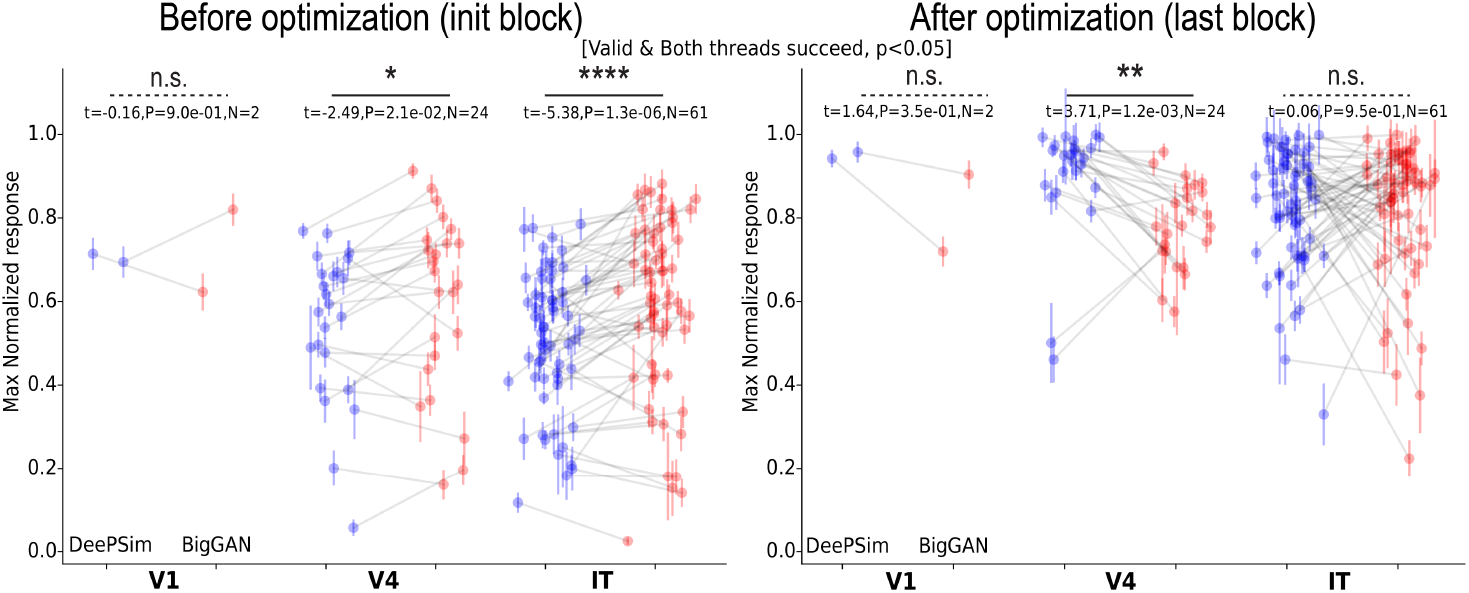
Neuronal activity rates before (left) and after optimization (right) *in vivo*. Activity rates were normalized by the maximum block-wise mean activation (across blocks and threads). Only experiments where both threads succeeded were included. Similar format as in Fig.3C.

**Fig. 9.**
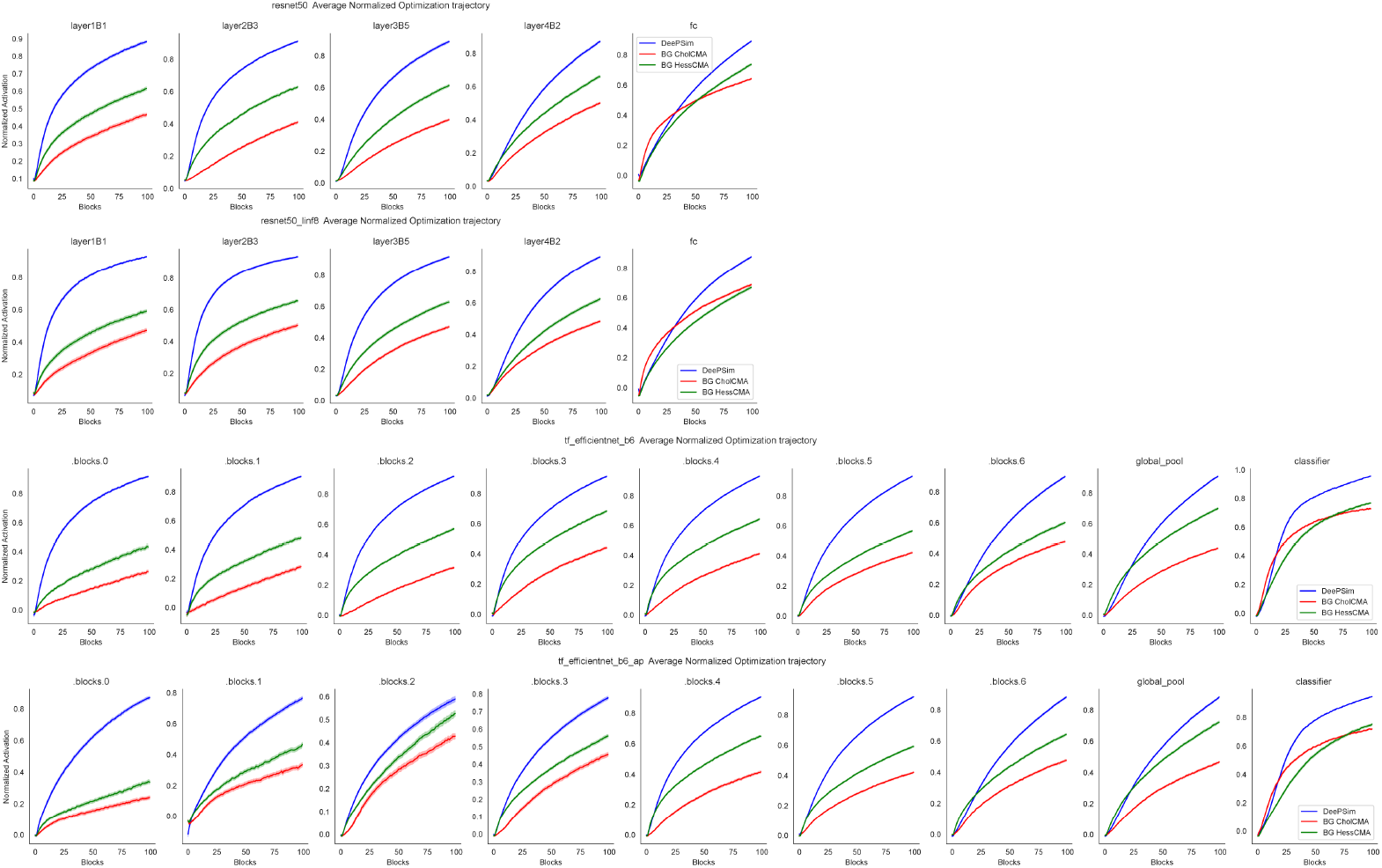
Evolution trajectory in DeePSim and BigGAN Space for Units *in silico*. Top to bottom: ResNet50, ResNet50-robust, EfficientNet-B6, EfficieintNet-B6 AdvProp, 50 units were sampled from each major layers, and 10 repeated evolutions were conducted in both DeePSim and BigGAN space. Consistently, driven by the same unit, DeePSim evolution reached higher activation than BigGAN, with the gap closer for deeper units in the network. Similar format as in Fig.4C.

**Fig. 10.**
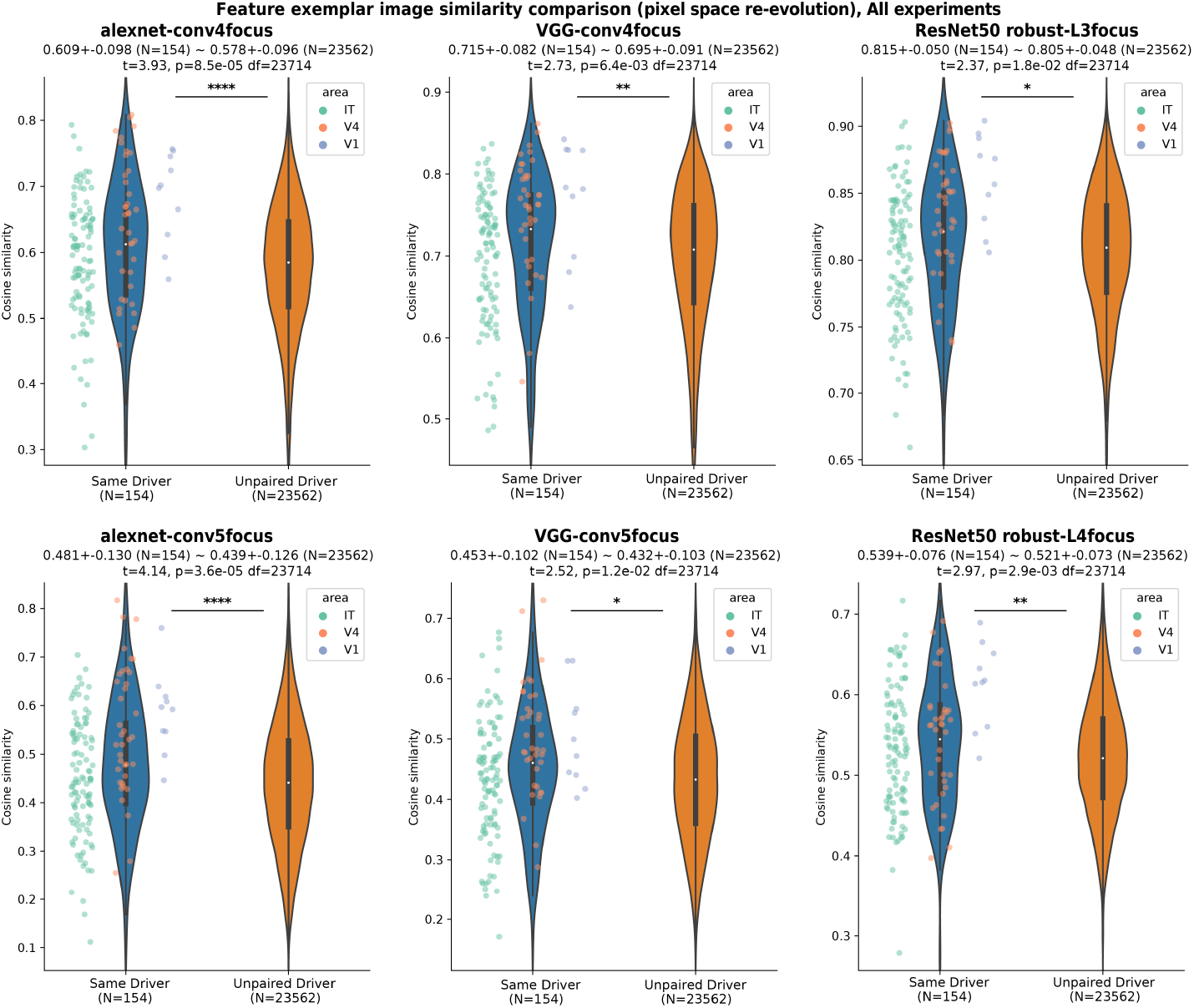
Similarity of prototypes in DeePSim and BigGAN space for units *in vivo* with various similarity metrics. Similar format as in Fig.6D, but with different image embedding spaces. From top to bottom, left to right: AlexNet conv4, conv5, VGG conv4, conv5, ResNet50-robust, layer3, layer4. For each layer, we averaged the activation tensor (weighted, along the spatial dimension) in that layer, to obtain an embedding vector with the same dimension as the channel dimension. The spatial averaging weight was a simple matrix that highlighted the center of the feature map and gradually fell off to zero at the border. This weight was a heuristic to emphasize features that were spatially more aligned with the neuronal receptive field. All 154 experiments were included. Consistently across metrics, feature exemplars from DeePSim and BigGAN space were more similar when they were driven by the same neuronal site than when driven by different sites.

**Fig. 11.**
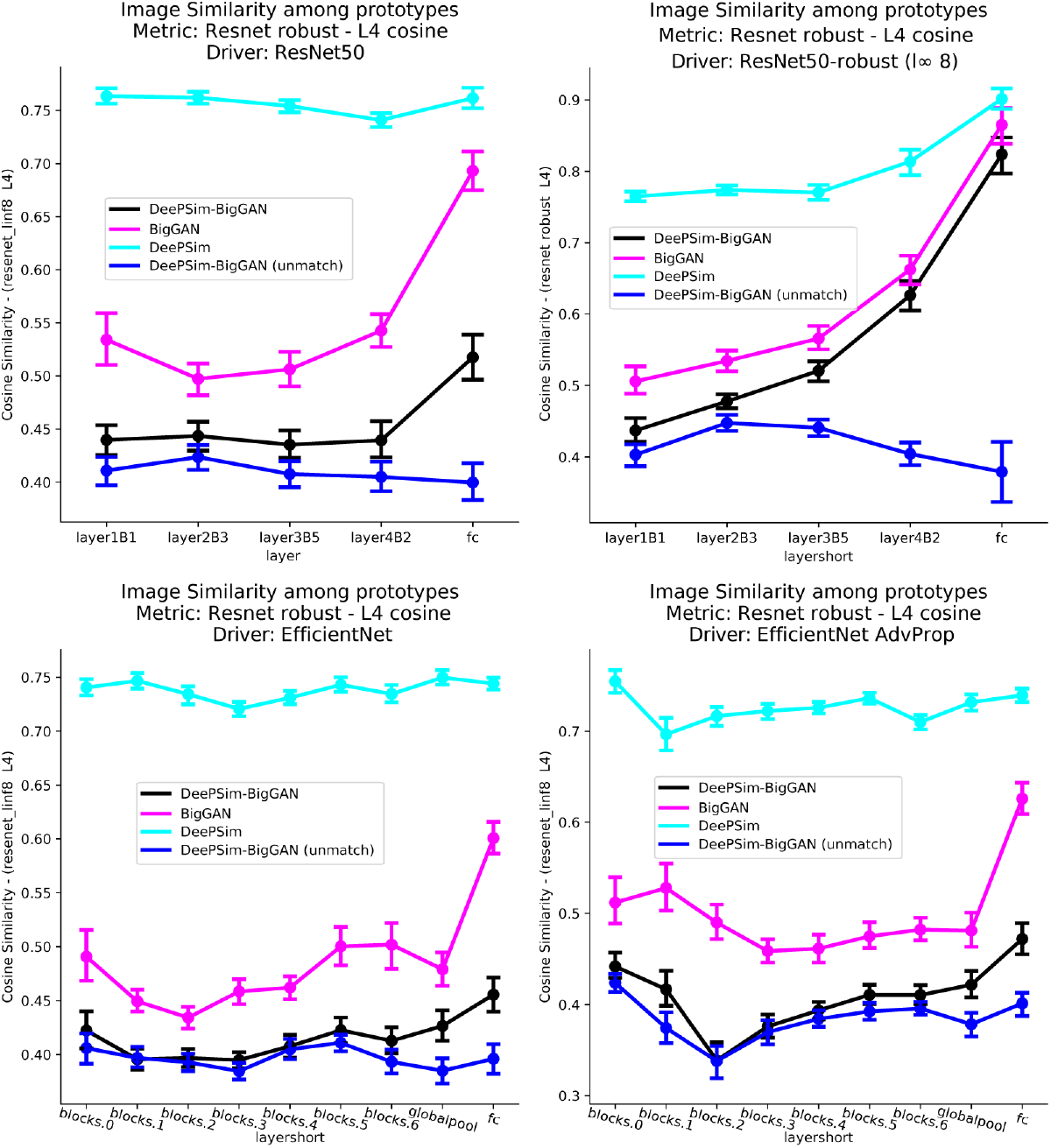
Similarity of Prototypes in DeePSim and BigGAN Space for Units *in silico*. Top to bottom, left to right: ResNet50, ResNet50-robust, EfficientNet-B6, EfficientNet-B6 AdvProp, 50 units were sampled from each major layer, and 10 repeated evolutions were conducted in both DeePSim and BigGAN space. Consistently, driven by the same unit, two prototypes from DeePSim space were more similar to one another than two prototypes in BigGAN space, than between DeePSim and BigGAN space. Prototypes of the same unit were more similar than ones of different units, especially for units deeper in the hierarchy. Similar format as in Fig.6E.

To measure local similarity, we developed a concentration index, a metric that measures the local concentration of activity by convolving the map with local Gaussian filters with diameters less than the full image. We first developed the concentration index *in silico*, by running parallel evolutions using AlexNet *conv5* units, and found that the local concentration index measured 0.0017±0.0002 (mean±SEM) between DeePSim-BigGAN images optimized by the same unit, vs. 0.0015±0.00008 between images optimized by different units (P = 7.4 × 10^−14^, Z-Val: 7.5, N = 91, Wilcoxon signed-rank test, Fig. 7E). Given the same analysis parameters, we then measured the mean concentration index values for optimized images driven by V1, V4, and IT neurons in monkeys A and B. We found that the mean concentration index for evolutions arising from the same neuronal drivers was 0.0017 ± 0.00010, and from different drivers, 0.0016 ± 0.00008 (P = 3.2 × 10^−8^, Z-Val: 5.5, N = 163, Wilcoxon signed-rank test). So, the DeePSim-BigGAN images were more likely to resemble each other when optimized by the same driving site, compared to images optimized by different sites, and this was due to local patch similarity.

## 2 Discussion

Most visual cortex neurons encode patterns that have never been hypothesized in previous studies, as shown by numerous closed-loop image synthesis studies [3, 4, 27, 28, 29, 30]. By leveraging generative networks that internalize natural image statistics, each representing image manifolds, we can identify these patterns and reveal the neuronal sites true activation range and tuning landscape [5]. However, how do these neuronal properties depend on the generative space? Here, we used two types of generators, specifically including one that creates images that are more photo-realistic and object-centric (BigGAN). We found an increasing alignment of neuronal tuning with the BigGAN object image manifold than the DeePSim image manifold along the ventral stream: V1 and V4 neurons aligned well with the more abstract, pattern based generator (DeePSim), whereas PIT neurons showed a comparable alignment to the DeePSim and BigGAN generators. When both BigGAN and DeePSim evolutions succeeded, V4 neurons showed an activity gap between optimized images, while the gap was closed for PIT neurons. Still, time mattered. The initial responses of PIT neurons were more aligned with the local-pattern DeePSim manifold, whereas later responses were more aligned with the configurational object (BigGAN) manifold. Overall, these results suggest that while the space of all possible naturalistic images is vast, and generative networks can only abstract limited portions of it, neurons in early ventral stream can respond strongly to every major manifold, suggesting that these networks truly encode local features also represented by the brain itself. Visual areas appear to align preferentially to different generative spaces. In principle, when the parameters that control a given image manifold are well-aligned with the “tuning parameters” encoded by visual areas, then the neurons’ tuning landscapes should be smoother, and easier for an evolutionary optimizer to navigate. In V4, the easier hill climbing in the DeePSim latent space versus the BigGAN latent space argues that the neurons’ tuning aligned more with the local, abstract axes in the DeePSim space than the BigGAN space; while in PIT, the alignment to two spaces seemed comparable at least, suggesting that both BigGAN and DeePSim have identified PIT-preferred features.

Although we have tested two different spaces based on generative networks, there are hundreds of alternatives in the existing literature alone. In theory, each visual neuron might be optimally aligned with a different image manifold, forming a unique energy landscape on the image space. It is crucial to test whether more anterior regions in the ventral stream align even more with the photo-realistic, object-based BigGAN image parametrization. However, one of the most important findings in this study was to show the importance of combining different generators when performing feature visualization, in order to prevent over-interpretation of results. While neuron-guided image optimization may lead to an object, that is not sufficient evidence to conclude that the neuron *encodes* that object, if one is using generative networks that randomly emit objects. A necessary control is to allow neuron-guided image optimization with generative networks that are not biased to create objects. Images containing objects and images lacking objects can be equally activating for a given neuron or neuronal microcluster in the ventral stream, which suggests that the underlying representation may be local and part of a distributed representation. This confound is a particular risk with the increasing use of modern generative models that assimilate natural images and art (e.g. diffusion models[31]), which might tempt experimental designs forcing perceptually aligned, semantic images onto neuronal functions. Using a spectrum of generative models parameterizing different image manifolds will provide a more accurate perspective on the latent factors that drive the visual system.

## 3. Methods

### 3.1 General Setup

Each experimental session was controlled using MonkeyLogic2[32], which directed the presentation of visual stimuli on ViewPixx EEG monitors (ViewPixx Technologies). The refresh rate was 120 Hz at a resolution of 1920 × 1080 pixels. The monkeys were located 58 cm from the screen. Gaze position was tracked via ISCAN cameras (ISCAN Inc.). Animals fixated 0.25°-diameter circles within a 1.0–2.0°-wide window during stimulus presentation; they obtained a reward if they held their gaze on the target for 2-3 s. Rewards are delivered via the DARIS Control Module System (Crist Instruments).

### 3.2 Research Animals

Two male adult monkeys (A and B, Macaca *mulatta*, aged 10 years, 10-11 kg) were implanted with chronic floating microelectrode arrays (Microprobes for Life Sciences, MD) in the right hemisphere: one array was located at the posterior lip of the lunate sulcus (corresponding to the V1/V2 transition), one on the prelunate gyrus (V4) and another anterior to the inferior occipital sulcus (PIT). We refer to these sites as V1/V2, V4 and PIT. The locations of the arrays were chosen based on sulcal landmarks, fine-tuned by local vasculature.

### 3.3 Neuronal Recordings

Neuronal electrical activity was acquired using OmniPlex Neural Recording Data Acquisition Systems (Plexon Inc.), with the PlexControl client to sort electrical events online based on waveform and inter-spike-intervals. Because all evolution experiments were based on a real-time, closed-loop design between neuronal activity and image synthesis, spike sorting was done manually at the beginning of each experiment. Within each channel, sources were estimated as arising from single-units, multi-units, or “hash” using a 1-5 scale, where 1 indicated strong confidence on the presence of a single-unit (based on waveform shape, inter-spike interval and separation from the main hash signal) and 5 reflecting multiunit activity that would not be separable. We use the term *site* to refer to all signal types; across experiments, sites comprised mostly multiunits/hash and some single units. After data collection, all spike/event times were discretized into 1-ms bins and convolved with a symmetric Gaussian probability density function with a 2-ms standard deviation.

### 3.4 Image Generative Models

For DeePSim, we used the fc6 model[7], with a custom Matlab implementation translated from the original `Caffe` model and weights. For BigGAN, we used the BigGAN-deep-256 version, implemented in `pytorch-pretrained-BigGAN`.

### 3.5 Image Statistics

To measure the FID score, we used the function `calculate_frechet_distance` from the library `pytorch-gan-metrics`.

The set of images used to calculate these statistics were

1. **ImageNet** 50000 images from the ImageNet validation set.
2. **DeePSim** 50000 images generated from DeePSim FC6 network, the latent codes were sampled from an isotropic Gaussian distribution with std. 4.
3. **BigGAN** 50000 images, 50 images per class for the 1000 classes. For each class, we used the pretrained object vector *c* for that class and 50 noise vectors *z* sampled from a 128d truncated Gaussian with truncation 0.7.
4. **BigGAN-RND** generated from BigGAN with 50000 latent vector sampled from 256d spherical Gaussian with std 0.08.

Additionally, we quantified low-level image feature values across both generators. First, we optimized images for units in AlexNet conv5 units (with receptive fields placed at the center, N = 91 units); a preferred image for each unit was concurrently optimized using the DeePSim- and the BigGAN generators, as described above. We used all images in the last evolved generation (N = 3640 total). Each image was analyzed as follows:

#### Luminance

We calculated the mean luminance of each image in a given set and constructed a histogram representation of the luminance distribution for each image. First, each image was converted from RGB to grayscale, then computed the global mean of the matrix elements. We also constructed a luminance histogram using the *imhist*.*m* function.

#### Contrast

First, each image was converted from RGB to grayscale. Then, the standard deviation of the pixel intensity values in the grayscale image was computed to represent the root-mean-square (RMS) contrast. The standard deviation calculation was performed using the MATLAB function *std*.*m* on the pixel intensity values of the grayscale image.

#### Spatial Frequency

We calculated the total power of the spatial frequency content and the edge density for each image in a given set. Each image was first converted from RGB to grayscale using the *rgb2gray*.*m* function. The 2D Fast Fourier Transform (FFT) of the grayscale image was then computed using the *fft2*.*m* function, and the power spectrum was obtained by taking the squared magnitude of the shifted Fourier-transformed image using *fftshift*.*m*. The total power of the power spectrum was calculated as the sum of all squared magnitudes. Edges were detected using the Canny method with the *edge*.*m* function, and the edge density was calculated as the proportion of edge pixels to the total number of pixels in the image.

#### Texture

Texture features were computed for each image in a given set using two methods: Gray Level Co-occurrence Matrix (GLCM) and Local Binary Patterns (LBP). Each image was first converted to grayscale using the *rgb2gray*.*m* function. The GLCM was computed using the *graycomatrix*.*m* function, and its properties—contrast, correlation, energy, and homogeneity—were extracted using the *graycoprops*.*m* function. Additionally, LBP features were extracted using the *extractLBPFeatures*.*m* function, providing a descriptor of the local spatial structure and contrast of the image. *Color*. Color distribution was computed for each image by measuring the histogram and balance for each color channel (red, green, and blue). For each image, the histogram of each color channel was computed using the *imhist*.*m* function. Additionally, the color balance for each channel was determined by calculating the mean intensity value.

#### Sharpness

We calculated the sharpness of each image in a given set by first converting the image to grayscale using the *rgb2gray*.*m* function if it was in color. We then used the Sobel operator to compute the gradient in both the X and Y directions, using convolution with the Sobel kernels defined in the *conv2*.*m* function. The gradient magnitudes were obtained by computing the square root of the sum of the squares of the gradients in the X and Y directions. Finally, the sharpness was quantified as the mean magnitude of these gradients, providing a measure of the image’s detail.

#### Shape Statistics

Other shape statistics were computed for each image by first converting the RGB images to grayscale. A binary image was then obtained through thresholding with the *graythresh*.*m* and *imbinarize*.*m* functions. The Euler number, representing the number of objects minus the number of holes, was calculated using the *bweuler*.*m* function. For each object in the binary image, we computed the aspect ratios of their bounding boxes (the ratio of width to height) using the *regionprops*.*m* function, focusing on the largest detected object by area.

#### Frequency Power Distribution

We calculated the frequency power distribution in vertical, horizontal, and diagonal directions for each image. Each image was first converted to grayscale, and we then applied the Fourier Transform and shifted the zero-frequency component to the center of the spectrum. The magnitude spectrum was obtained by taking the absolute value of the shifted Fourier Transform. We calculated the overall power as the sum of all magnitudes. The vertical power was computed as the sum of magnitudes along the vertical central line, the horizontal power as the sum of magnitudes along the horizontal central line, and the diagonal power as the sum of magnitudes along the diagonal line from the top-left to the bottom-right, each normalized by the overall power.

#### Symmetry

The symmetry of each image was calculated across the horizontal, vertical, and radial axes. Each image was first converted to grayscale. Horizontal symmetry was assessed by comparing the top half of the image to the flipped bottom half, calculating the mean absolute difference between corresponding pixels. Vertical symmetry was evaluated similarly by comparing the left half to the flipped right half. Radial symmetry was determined by creating concentric rings of pixels around the center of the image and comparing each pixel in the ring to the mean pixel value of that ring, with the radial symmetry score being the average absolute difference from the mean.

#### Entropy

The Shannon entropy and joint entropy for each image were also computed. The Shannon entropy, which measures the information content of the grayscale representation, was computed by first converting each RGB image to grayscale and then applying the *entropy*.*m* function. The joint entropy, which measures the combined information content of the RGB channels, was computed by flattening the RGB channels and creating a joint histogram using the *imhist*.*m* function. The joint entropy was then calculated from this histogram by normalizing the counts and summing the product of the non-zero histogram values with their log base 2 values.

### 3.6 Evolutionary algorithm

We used and adapted the evolutionary algorithms developed in our previous studies [8, 9]. For the DeePSim evolutions, we used the *Cholesky* covariance matrix adaptation evolutionary strategy (CMA-ES) algorithm[33] and the *HessianCMA* algorithm. The Cholesky CMAES has the same parameters as in previous works [4, 5]: standard deviation was initialized as 3.0. For the *HessianCMA* Optimizer, we pre-computed the 500 most informative dimensions in the latent space of the DeePSim GAN through Hessian decomposition [8], and only optimized in that linear subspace, which was more sample efficient and we have shown it yielded comparable activation with the optimizer operating in the full 4096D space. For the BigGAN evolutions, we used the Cholesky CMAES and *HessianCMA* algorithms [8] with the optimized parameters optimized on *in silico* experiments, primarily choosing an initial standard deviation of 0.06.

### 3.7 Neuron-guided image synthesis (“evolutions”)

We used the same experimental protocol for neuron-guided image synthesis as in previous studies[5]. In each paired-evolution experiment, first, one neuronal site was chosen as a target site (driver). Driver units were selected if they had a well-defined receptive field and visually evoked responses relative to baseline. We first determined the optimal location (image center and size) for the driver using RF mapping experiments. Then we placed the stimuli at the optimal location and started the Paired Evolution experiments. We have two image generators; each has a set of 30 initial images as the starting point. For DeePSim generator these initial images were Portilla and Simoncelli textures[34], inverted into the DeePSim space. The original embedding position or image identity of the first generation was not crucial, but these inverted texture images were used because localized the first generation to the origin of the latent space (input vectors had very small norms). For the BigGAN generator, initial images were generated from random vectors, also having small vector norms. Each experiment started by presenting these initial images along with 10 reference, static images. Reference images were selected if they were known to evoke high activity from the array site under study, per prior experiments. After all images were presented once, the driver’s spike-rate responses during 50-200 ms post image onset was averaged as the score of the corresponding image. The latent vectors from each generator and their corresponding scores were provided to the evolutionary optimizer as tuned for this generative space. Then, the optimizer returned the next batch of latent vectors for the corresponding generator. These vectors were rendered by the generator to create new images. Finally, the new batch of images from both generators were randomly interleaved into a sequence and presented to the monkey, starting the new block or iteration. Each experiment comprised 10 to 60 blocks, and was stopped after the neuronal firing rate converged or stopped increasing for both threads. Each image was presented once. The same procedure was carried out when optimizing the activity of CNN units.

### 3.8 Alignment and average optimization trajectory

We computed the average optimization trajectory across sessions. Because different sessions had different numbers of blocks over the evolution, we lined up all experiments to the same x-axis by extrapolating the evolution trajectory by the mean activation of the last 2 blocks, to match the number of blocks of the longest session (56 blocks). We also kept the variability within those last 2 blocks. Because different neurons and neuronal microclusters have different ranges of activity, for each session, we first normalized the neuronal activation in each block by the maximum block-averaged activation of that driver unit across the two spaces. Then we pooled across sessions and averaged the length-aligned optimization trajectories.

### 3.9 Quantifying evolution convergence speed

To quantify the speed at which neurons and microclusters achieved their maximum activity during the evolution (convergence speed), we fit a Gaussian process regression model for the optimization trajectory: neuronal activation as a function of block number. Then, for the smoother GP regressed curve, we identified the number of blocks before the trajectory reached 80% maximal activation for that thread. While there were other ways to compute the time-constant statistics, we found this choice did not affect the trend (Tab. 4).

### 3.10 Similarity of peristimulus time histograms

To quantify the distance between paired-evolution PSTHs, we used two measures: one integrated the *absolute* area between the difference of the two curves (Eq. 1), and the other computed the difference between the averaged level of PSTH curve, equivalently integrating the *signed* area between the PSTH curves (Eq. 2).

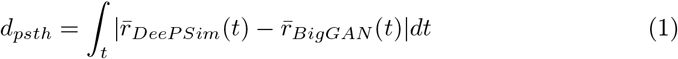

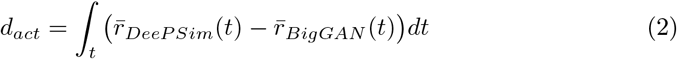

For both measures, we first normalized the PSTHs, i.e. divided them by the max block-average firing rate for that neuron.

### 3.11 Similarity of images

Image similarity was computed in different feature spaces. For the first image analysis, we chose ResNet50-robust as our image embedding model and computed the cosine similarity of each image vector embedding as the similarity score. For the second analysis, we used the Learned Perceptual Image Patch Similarity (LPIPS) metric [25]. The LPIPS object class was instantiated with the AlexNet backbone. The LPIPS object returns a spatial map of perceptual distances the same size as image inputs. This procedure was first tested using CNN-unit-driven evolutions, specifically AlexNet convolutional layer 5 (N = 91 randomly sampled units). For each paired-evolution experiment (given one CNN unit), we sampled 15 images from the final generation of DeePSim images and 15 images from the final generation of BigGAN images, then compared each of the DeePSim images to the other 15 BigGAN images (we also used BigGAN as references with the same results). The core of our analysis involved calculating the perceptual similarity between each image pair. After we obtained the LPIPS distance map, we converted it into a similarity heatmap by subtracting it from 1 (similarity = 1 – LPIPS ). A key part of our analysis involved quantifying the concentration of perceptual similarity within the heatmaps. To achieve this, we calculated a *concentration score* from a given heatmap. The concentration score served as a quantitative measure of the degree to which perceptual similarity was localized within specific regions of the heatmap. The concentration index function defined a range of filter sizes to be applied to the heatmap, starting from a minimum size (set to 1 for individual pixel consideration), with multiple ranges, allowing the function to assess concentration at various spatial scales. For each filter size within the specified range, the function created a uniform filter (a matrix of ones) of that size. This filter was then normalized to ensure that its total sum was 1, maintaining the scale of the original heatmap values. The normalized filter was convolved with the heatmap, effectively averaging the heatmap values over areas corresponding to the filter size. This convolution process was repeated for each filter size in the range. After each convolution, the maximum value from the convolution result was extracted and stored. These maximum values represented the highest concentration of perceptual similarity for each filter size, indicating the presence of localized high-similarity regions within the heatmap. The concentration score was computed as the mean of these maximum values. This score provided a single, comprehensive metric indicating the extent of localized perceptual similarity in the heatmap. A higher concentration score suggested more pronounced localization of similarity, while a lower score indicated more diffuse similarity across the image. Next, for each reference DeePSim image for a given unit *u*_*i*_, we randomly sampled another experiment and compared the unit *u*_*i*_-generated DeePSim image to 15 of the random unit *u*_*j*_-generated BigGAN images. We then compared the average similarity between the same-driver image pairs (*u*_*i*_-generated DeePSim image vs. *u*_*i*_-generated BigGAN image) to the average similarity between the different-driver image pairs (*u*_*i*_-generated DeePSim image vs. *u*_*j*_-generated BigGAN image), and tested for statistical significance using a Wilcoxon signed-rank test.

### 3.12 Feature attribution of evolved images

To determine and emphasize image features that were important during the evolution, we used a previously published method [26]. We used every pair of generated image and its associated neuronal response during a given evolution to build an encoding model. We found all the model units that correlated with the given neuronal firing rate. Then we factorized the correlation matrix, and used it as an initialization for weights, we optimized this factorized read-out weight matrix similar to [35]. We then used gradient-based optimization to find the maximum firing rate stimulus for the model unit: the optimized image was called *feature exemplar* as showed in Fig.6C.

### 3.13 General Statistical Procedure

Neuronal sampling. We implanted arrays in cortical regions marked solely by sulcal patterns, without pre-selection via functional imaging, and as such, we achieved a random sample. Cortical sites were studied if they were visually responsive, as it is expected that some microwires could land outside of the cortical band and/or cause micro-infarcts. Our dataset included 87 unique electrode locations or neuronal sites. Most of these sites (51) were sampled once, while the rest were sampled multiple times: 21 sites twice, 6 sites three times, 4 sites six times, 2 sites five times, and 1 site nine times. These sites show some correlation over time, although they cannot be designated as identical neurons or neuronal populations.

We used the Student’s t-test to perform two-sample tests of statistical significance (which assumes normality). Reported P-values were calculated using two-sided tests, unless otherwise stated.

## Data availability

Processed data for the analysis and reproduction of the figures can be downloaded at https://osf.io/pre96. The complete raw dataset used in this study is available upon request to B.W. and C.R.P.

## Code availability

The code used to analyze the data in this manuscript will be made available on GitHub at https://github.com/Animadversio/Dynamic-Neuron-GAN-Alignment

## 4 Supplementary information

## Abbreviations

raw act: raw firing rate in 50-200ms window
evoke: raw firing rate subtracting a session averaged baseline firing rate computed from 0-40ms window;
bsl init: raw firing rate subtracting the mean firing rate in the initial block, namely activation increase from the initial block. We reported the
bsl init: in the main text and figure.

## References

[1] Ian Goodfellow, Jean Pouget-Abadie, Mehdi Mirza, Bing Xu, David Warde-Farley, Sherjil Ozair, Aaron Courville, and Yoshua Bengio. Generative adversarial nets. In Advances in Neural Information Processing Systems, pages 2672–2680, 2014.

[2] Tero Karras, Timo Aila, Samuli Laine, and Jaakko Lehtinen. Progressive growing of gans for improved quality, stability, and variation. In International Conference on Learning Representations, 2018.

[3] Carlos R. Ponce, Will Xiao, Peter F. Schade, Till S. Hartmann, Gabriel Kreiman, and Margaret S. Livingstone. Evolving Images for Visual Neurons Using a Deep Generative Network Reveals Coding Principles and Neuronal Preferences. Cell, 177(4):999–1009.e10, may 2019.

[4] Olivia Rose, James Johnson, Binxu Wang, and Carlos R Ponce. Visual prototypes in the ventral stream are attuned to complexity and gaze behavior. Nature communications, 12(1):1–16, 2021.

[5] Binxu Wang and Carlos R. Ponce. Tuning landscapes of the ventral stream. Cell Reports, nov 2022.

[6] Will Xiao and Gabriel Kreiman. XDream: Finding preferred stimuli for visual neurons using generative networks and gradient-free optimization. PLOS Computational Biology, 16(6):e1007973, jun 2020.

[7] Alexey Dosovitskiy and Thomas Brox. Generating images with perceptual similarity metrics based on deep networks. Advances in neural information processing systems, 29, 2016.

[8] Binxu Wang and Carlos R Ponce. A geometric analysis of deep generative image models and its applications. In International Conference on Learning Representations, 2020.

[9] Binxu Wang and Carlos R. Ponce. High-performance evolutionary algorithms for online neuron control. In Proceedings of the Genetic and Evolutionary Computation Conference, GECCO ‘22, page 1308–1316, New York, NY, USA, 2022. Association for Computing Machinery.

[10] Andrew Brock, Jeff Donahue, and Karen Simonyan. Large Scale GAN Training for High Fidelity Natural Image Synthesis. 7th International Conference on Learning Representations, ICLR 2019, sep 2018.

[11] Jia Deng, Wei Dong, Richard Socher, Li-Jia Li, Kai Li, and Li Fei-Fei. Imagenet: A large-scale hierarchical image database. In 2009 IEEE conference on computer vision and pattern recognition, pages 248–255. Ieee, 2009.

[12] Ting Chen, Mario Lucic, Neil Houlsby, and Sylvain Gelly. On self modulation for generative adversarial networks. arXiv preprint 1810.01365, 2018.

[13] Han Zhang, Ian Goodfellow, Dimitris Metaxas, and Augustus Odena. Selfattention generative adversarial networks. In International conference on machine learning, pages 7354–7363. PMLR, 2019.

[14] Martin Heusel, Hubert Ramsauer, Thomas Unterthiner, Bernhard Nessler, and Sepp Hochreiter. Gans trained by a two time-scale update rule converge to a local nash equilibrium. Advances in neural information processing systems, 30, 2017.

[15] Fu Xing Long, Bas van Stein, Moritz Frenzel, Peter Krause, Markus Gitterle, and Thomas Bäck. Learning the characteristics of engineering optimization problems with applications in automotive crash. In Proceedings of the Genetic and Evolutionary Computation Conference, pages 1227–1236, 2022.

[16] Daniel LK Yamins, Ha Hong, Charles F Cadieu, Ethan A Solomon, Darren Seibert, and James J DiCarlo. Performance-optimized hierarchical models predict neural responses in higher visual cortex. Proceedings of the national academy of sciences, 111(23):8619–8624, 2014.

[17] Simona Celebrini, Simon Thorpe, Yves Trotter, and Michel Imbert. Dynamics of orientation coding in area v1 of the awake primate. Visual neuroscience, 10(5):811–825, 1993.

[18] Dario L Ringach, Michael J Hawken, and Robert Shapley. Dynamics of orientation tuning in macaque primary visual cortex. Nature, 387(6630):281–284, 1997.

[19] Yasuko Sugase, Shigeru Yamane, Shoogo Ueno, and Kenji Kawano. Global and fine information coded by single neurons in the temporal visual cortex. Nature, 400(6747):869–873, 1999.

[20] Scott L Brincat and Charles E Connor. Dynamic shape synthesis in posterior inferotemporal cortex. Neuron, 49(1):17–24, 2006.

[21] Victor AF Lamme, Valia Rodriguez-Rodriguez, and Henk Spekreijse. Separate processing dynamics for texture elements, boundaries and surfaces in primary visual cortex of the macaque monkey. Cerebral cortex, 9(4):406–413, 1999.

[22] Victor AF Lamme and Pieter R Roelfsema. The distinct modes of vision offered by feedforward and recurrent processing. Trends in neurosciences, 23(11):571–579, 2000.

[23] Kaiming He, Xiangyu Zhang, Shaoqing Ren, and Jian Sun. Deep residual learning for image recognition. In Proceedings of the IEEE conference on computer vision and pattern recognition, pages 770–778, 2016.

[24] Aleksander Madry, Aleksandar Makelov, Ludwig Schmidt, Dimitris Tsipras, and Adrian Vladu. Towards deep learning models resistant to adversarial attacks. In International Conference on Learning Representations, 2018.

[25] Richard Zhang, Phillip Isola, Alexei A Efros, Eli Shechtman, and Oliver Wang. The unreasonable effectiveness of deep features as a perceptual metric. In Proceedings of the IEEE conference on computer vision and pattern recognition, pages 586–595, 2018.

[26] Binxu Wang and Carlos R Ponce. Factorized convolution models for interpreting neuron-guided images synthesis. 2022 Conference on Cognitive Computational Neuroscience, aug 2022.

[27] Chong Guo, Michael Lee, Guillaume Leclerc, Joel Dapello, Yug Rao, Aleksander Madry, and James Dicarlo. Adversarially trained neural representations are already as robust as biological neural representations. In International Conference on Machine Learning, pages 8072–8081. PMLR, 2022.

[28] Edgar Y Walker, Fabian H Sinz, Erick Cobos, Taliah Muhammad, Emmanouil Froudarakis, Paul G Fahey, Alexander S Ecker, Jacob Reimer, Xaq Pitkow, and Andreas S Tolias. Inception loops discover what excites neurons most using deep predictive models. Nature neuroscience, 22(12):2060–2065, 2019.

[29] Konstantin F Willeke, Kelli Restivo, Katrin Franke, Arne F Nix, Santiago A Cadena, Tori Shinn, Cate Nealley, Gabby Rodriguez, Saumil Patel, Alexander S Ecker, et al. Deep learning-driven characterization of single cell tuning in primate visual area v4 unveils topological organization. bioRxiv, pages 2023–05, 2023.

[30] Zhiwei Ding, Dat T Tran, Kayla Ponder, Erick Cobos, Zhuokun Ding, Paul G Fahey, Eric Wang, Taliah Muhammad, Jiakun Fu, Santiago A Cadena, et al. Bipartite invariance in mouse primary visual cortex. bioRxiv, 2023.

[31] Cody Nash. Can ai create a white painting?, 2023. Accessed: 2024-04-10.

[32] Jaewon Hwang, Andrew R. Mitz, and Elisabeth A. Murray. Nimh monkeylogic: Behavioral control and data acquisition in matlab. Journal of Neuroscience Methods, 323:13–21, 2019.

[33] Nikolaus Hansen and Andreas Ostermeier. Completely derandomized self-adaptation in evolution strategies. Evolutionary computation, 9(2):159–195, 2001.

[34] Javier Portilla and Eero P Simoncelli. A parametric texture model based on joint statistics of complex wavelet coefficients. International journal of computer vision, 40:49–70, 2000.

[35] David Klindt, Alexander S Ecker, Thomas Euler, and Matthias Bethge. Neural system identification for large populations separating “what” and “where”. Advances in neural information processing systems, 30, 2017.

